# *Caenorhabditis elegans* processes sensory information to choose between freeloading and self-defense strategies

**DOI:** 10.1101/2020.02.24.963637

**Authors:** Jodie Schiffer, Francesco Servello, William Heath, Francis Raj Gandhi Amrit, Stephanie Stumbur, Sean Johnsen, Julian Stanley, Hannah Tam, Sarah Brennan, Natalie McGowan, Abigail Vogelaar, Yuyan Xu, William Serkin, Arjumand Ghazi, Javier Apfeld

## Abstract

Hydrogen peroxide is the preeminent chemical weapon that organisms use for combat. Individual cells rely on conserved defenses to prevent and repair peroxide-induced damage, but whether similar defenses might be coordinated across cells in animals remains poorly understood. Here, we identify a neuronal circuit in the nematode *Caenorhabditis elegans* that processes information perceived by two sensory neurons to control the induction of hydrogen-peroxide defenses in the organism. We found that catalases produced by *Escherichia coli*, the nematode’s food source, can deplete hydrogen peroxide from the local environment and thereby protect the nematodes. In the presence of *E. coli*, the nematode’s neurons signal via TGFβ-insulin/IGF1 relay to target tissues to repress expression of catalases and other hydrogen-peroxide defenses. This adaptive strategy is the first example of a multicellular organism modulating its defenses when it expects to freeload from the protection provided by molecularly orthologous defenses from another species.

## Introduction

Bacteria, fungi, plants, and animal cells have long been known to excrete large quantities of hydrogen peroxide to attack their prey and pathogens (Avery and Morgan, 1924). Hydrogen peroxide is also a dangerous byproduct of aerobic respiration (Chance et al., 1979). Cells rely on highly conserved defense mechanisms to degrade hydrogen peroxide and avoid the damage that hydrogen peroxide inflicts on their proteins, nucleic acids, and lipids (Mishra and Imlay, 2012). The extent to which these protective defenses are coordinated across cells in animals is poorly understood.

In the present study, we used the nematode *C. elegans* as a model system to explore whether hydrogen-peroxide protective defenses are coordinated across cells. We focused on the role of sensory neurons in this coordination because sensory circuits in the brain collect and integrate information from the environment, enabling animals to respond to environmental change. Specific sensory neurons enable nematodes to smell, taste, touch, and sense temperature and oxygen levels (Bargmann and Horvitz, 1991a; Chalfie et al., 1985; Gray et al., 2004; Mori and Ohshima, 1995; White et al., 1986). This information is integrated rapidly by interneurons to direct the nematode’s movement towards favorable environmental cues and away from harmful ones (Kaplan et al., 2018). Nematodes also use sensory information to modify their development, metabolism, lifespan, and heat defenses (Apfeld and Kenyon, 1999; Bargmann and Horvitz, 1991b; Mak et al., 2006; Prahlad et al., 2008). Understanding how sensory circuits in the brain regulate hydrogen-peroxide defenses in *C. elegans* may provide a template for understanding how complex animals coordinate their cellular defenses in response to the perceived threat of hydrogen-peroxide attack.

Using a systematic neuron-specific genetic-ablation approach, we identified ten classes of sensory neurons that regulate sensitivity to harmful peroxides in *C. elegans*. We found that the two ASI sensory neurons of the amphid, the major sensory organ of the nematode, initiate a multistep hormonal relay that decreases the nematode’s hydrogen peroxide defenses: a DAF-7/TGFβ signal from ASI is received by multiple sets of interneurons, which independently process this information and then relay it to target tissues via insulin/IGF1 signals. Interestingly, this neuronal circuit lowers the action of endogenous catalases and other hydrogen-peroxide defenses within the worm in response to perception and ingestion of *E. coli*, the nematode’s primary food source in laboratory experiments. We show that *E. coli* express orthologous defenses that degrade hydrogen peroxide in the environment and that *C. elegans* does not need to induce catalases and other hydrogen-peroxide defenses when *E. coli* is abundant. Thus, this neuronal circuit enables the nematodes to lower their own defenses upon sensing bacteria that can provide protection. In the microbial battlefield, nematodes use a sensory-neuronal circuit to determine whether to defend themselves from hydrogen peroxide attack or to freeload off protective defenses from another species.

## Results

### Sensory neurons regulate peroxide resistance in *C. elegans*

*C. elegans* is highly sensitive to the lethal effects of peroxides. Under standard laboratory conditions, wild-type nematodes have an average lifespan of approximately 15 days (Kenyon et al., 1993). In contrast, when grown in the presence of peroxides (6 mM tert-butyl hydroperoxide, tBuOOH), the average lifespan of these nematodes is reduced to less than 1 day (Fig. 1A) (An et al., 2005). Previously, we determined the peroxide resistance of nematodes by measuring their lifespan with high temporal resolution in the presence of 6 mM tBuOOH (Stroustrup et al., 2013).

**Figure 1.**
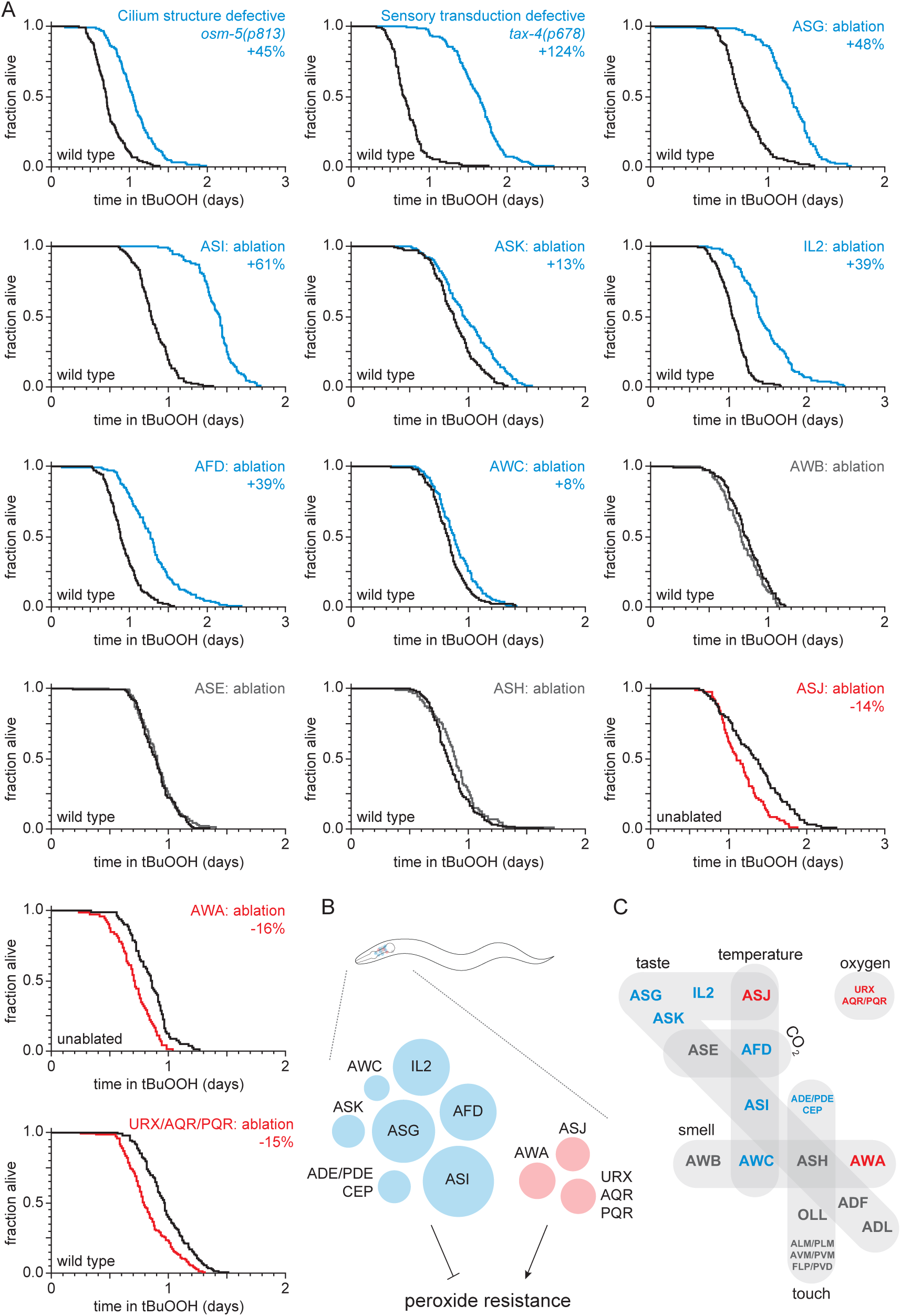
Sensory neurons regulate peroxide resistance in *C. elegans*. (A) Peroxide resistance of nematodes with defects in sensory cilia and sensory transduction, or with genetic ablation of specific sensory neurons. The fraction of nematodes remaining alive in the presence of 6 mM tert-butyl hydroperoxide (tBuOOH) is plotted against time. Interventions that increased, decreased, or did not affect survival are denoted in blue, red, and gray, respectively, and their effects on mean peroxide resistance are noted. (B) Specific sensory neurons normally reduce (blue) or increase (red) peroxide resistance. Circle area denotes the effect of ablation of the respective neuron on mean peroxide resistance. (C) Sensory neurons are grouped by the stimuli they sense. Neurons that normally reduce (eight classes) or increase (three classes) peroxide resistance are shown in blue and red, respectively. See also Figure S1. Statistical analyses are in Table S1.

To investigate whether sensory neurons might regulate the nematode’s peroxide defenses, we measured peroxide resistance in mutant animals with global defects in sensory perception. We first examined *osm-5* cilium structure mutants, which lack neuronal sensory perception due to defects in the sensory endings (cilia) of most sensory neurons (Perkins et al., 1986). These mutants exhibited a 45% increase in peroxide resistance relative to wild-type controls (Fig. 1A and Table S1). Next, we examined *tax-2* and *tax-4* cyclic GMP-gated channel mutants, which are defective in the transduction of several sensory processes including smell, taste, oxygen, and temperature sensation (Coburn and Bargmann, 1996; Komatsu et al., 1996). These two mutants also exhibited large increases in peroxide resistance compared to wild-type controls (Figs. 1A and S1A, and Table S1). Together, these observations indicate that neuronal sensory perception plays a role in regulating peroxide resistance in nematodes.

In *C. elegans* hermaphrodites, 60 ciliated and 12 non-ciliated neurons perform most sensory functions (White et al., 1986). To identify which of these sensory neurons influence the nematode’s peroxide resistance, we systematically measured peroxide resistance in a collection of strains in which specific sensory neurons have been genetically ablated via neuron-specific expression of caspases (Chelur and Chalfie, 2007) or, in one case, via mutation of a neuron-specific fate determinant (Chang et al., 2003; Uchida et al., 2003). Overall, our neuron-ablation collection covered 44 ciliated and 10 non-ciliated neurons, including each of the 12 pairs of ciliated neurons that make up the two amphids (the major sensory organs), 8 of the 13 classes of non-amphid ciliated neurons, and 6 of the 7 classes of non-ciliated sensory neurons (Table S9). Individual ablation of ASI, ASG, ASK, AFD, AWC, IL2 and joint ablation of ADE, PDE, and CEP increased the nematode’s peroxide resistance by up to 61% (Figs. 1A-B and S1B-C, and Table S1), whereas individual ablation of ASJ and AWA, and joint ablation of URX, AQR, and PQR reduced peroxide resistance by up to 16% (Fig. 1A-B). The remainder of the neurons tested—ADF, ADL, ASE, ASH, AWB, OLL, and joint ablation of ALM, PLM, AVM, PVM, FLP, and PVD—did not affect peroxide resistance (Figs. 1A, 1C and S1D-G, and Table S1). Altogether, we found that ten classes of sensory neurons can positively or negatively modulate peroxide resistance (Fig. 1B). These neurons are known to respond to diverse stimuli, including smell, taste, touch, temperature, and oxygen levels (Fig. 1C), suggesting that nematodes might adjust their peroxide resistance in response to multiple types of sensory information.

### ASI sensory neurons regulate peroxide resistance via DAF-7/TGFβ signaling

Among all neuronal ablations tested, ablation of ASI, a pair of neurons that sense taste and temperature, caused the largest increase in peroxide resistance (Fig. 1A). Thus, we focused on the role of the ASI neuronal pair. ASI neurons secrete many peptide hormones, including DAF-7 (Meisel et al., 2014; Ren et al., 1996), a transforming growth factor β (TGFβ) hormone that regulates feeding, development, metabolism, and lifespan (Dalfo et al., 2012; Greer et al., 2008; Ren et al., 1996; Shaw et al., 2007). To determine whether DAF-7/TGFβ signaling also regulates peroxide resistance, we examined mutants in *daf-7*. We found that *daf-7(ok3125)* null and *daf-7(e1372)* loss-of-function mutations increased peroxide resistance two-fold relative to wild-type controls (Figs. 2A, 2B, and S2A-D, and Table S2). Reintroducing the *daf-7(+)* gene into *daf-7(ok3125)* mutants restored peroxide resistance to wild-type levels (Fig. 2B and Table S2). Moreover, expression of *daf-7(+)* only in the ASI neurons was sufficient to reduce the peroxide resistance of *daf-7(ok3125)* mutants to wild-type levels (Fig. 2C and Table S2). *daf-7* is also expressed at a low level in ASJ, another pair of chemosensory neurons (Meisel et al., 2014), and expression of *daf-7(+)* only in ASJ rescued the increased peroxide resistance of *daf-7(ok3125)* mutants (Fig. 2D and Table S2). Thus, expression of *daf-7* in ASI or ASJ was sufficient to confer normal peroxide resistance. Because ablation of ASI increased peroxide resistance but ablation of ASJ did not (Fig. 1A), we reason that ASI neurons are the source of DAF-7 that regulates the nematode’s peroxide resistance.

**Figure 2.**
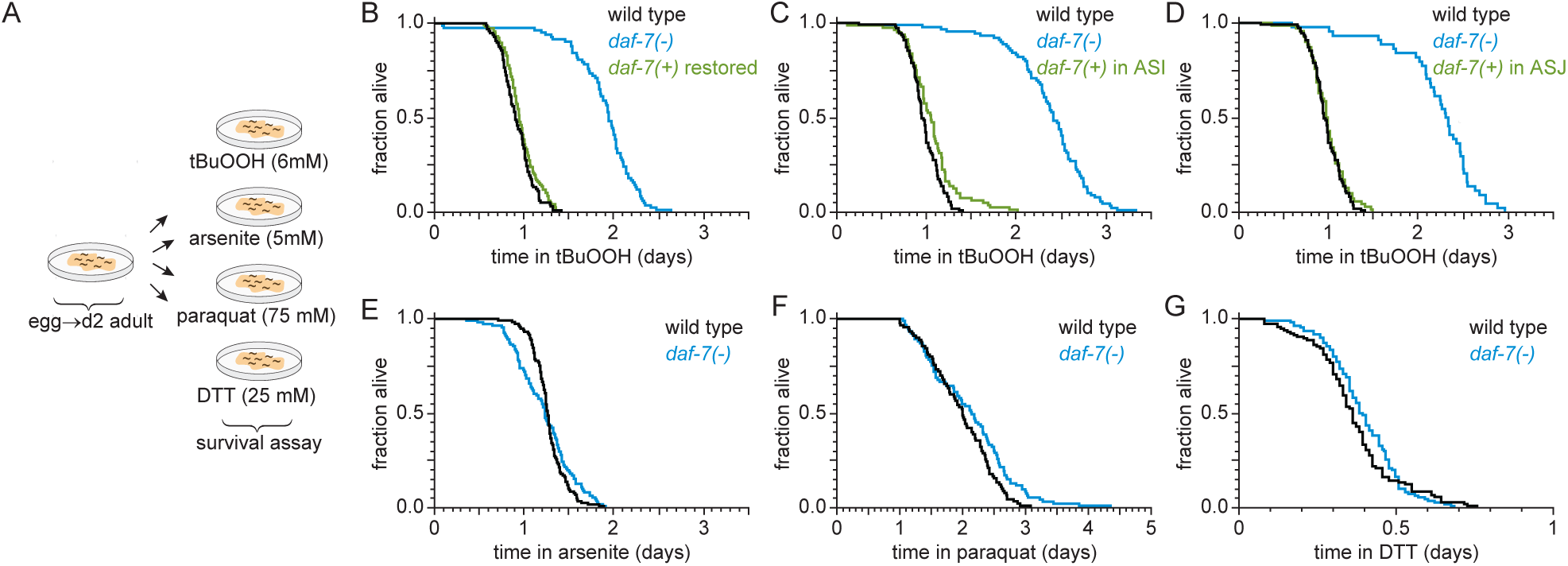
ASI sensory neurons secrete DAF-7/TGFβ to specifically lower the nematode’s peroxide resistance. (A) Diagram summarizing experimental strategy. (B-D) Peroxide resistance of wild type, *daf-7(ok3125)*, and *daf-7(ok3125)* with *daf-7(+)* reintroduced with (B) its endogenous promoter, (C) the ASI-specific *str-3* promoter, or (D) the ASJ-specific *trx-1* promoter. (E-G) Resistance to 5 mM arsenite, 75 mM paraquat, and 25 mM dithiothreitol (DTT) of wild type and *daf-7(ok3125)*. See also Figure S2. Statistical analyses are in Table S2.

We next asked whether DAF-7/TGFβ from ASI might regulate resistance to additional toxic chemicals from the environment that are not peroxides or directly generate peroxides. We tested sensitivity of *daf-7* mutants to arsenite (a toxic metalloid), paraquat (a redox-cycling herbicide), and dithiothreitol/DTT (a reducing agent). Compared with wild-type animals, *daf-7(ok3125)* mutants had similar survival in 5 mM arsenite, 25 mM dithiothreitol, and 75 mM paraquat (Figs. 2A and 2E-G, and Table S2). Therefore, the DAF-7/TGFβ signal from ASI is a specific regulator of peroxide resistance in the worm.

### Interneurons must reach a consensus to increase peroxide resistance in response to DAF-7/TGFβ from ASI

DAF-7/TGFβ signals via the Type 1 TGFβ receptor DAF-1 (Georgi et al., 1990) to regulate multiple downstream processes (Dalfo et al., 2012; Greer et al., 2008; Ren et al., 1996; Shaw et al., 2007). Signaling through the DAF-1 receptor inactivates the transcriptional activity of a complex between the receptor-associated coSMAD, DAF-3, and the Sno/Ski factor, DAF-5 (da Graca et al., 2004; Patterson et al., 1997; Tewari et al., 2004). We found that a similar signal-transduction pathway regulates peroxide resistance. *daf-1(m40)* loss-of-function mutants showed a two-fold increase in peroxide resistance (Fig. 3A and Table S3), and the increase in peroxide resistance of *daf-7* and *daf-1* mutants was almost completely abrogated by null or loss-of-function mutations in either *daf-3* or *daf-5* (Figs. 3A and S3A-C, and Table S3). The *daf-3(mgDf90)* null mutation also suppressed the increase in peroxide resistance of ASI-ablated worms (Fig. 3B and Table S3). Therefore, the ASI neurons normally function to lower peroxide resistance in the worm using a canonical TFGβ signaling pathway.

**Figure 3.**
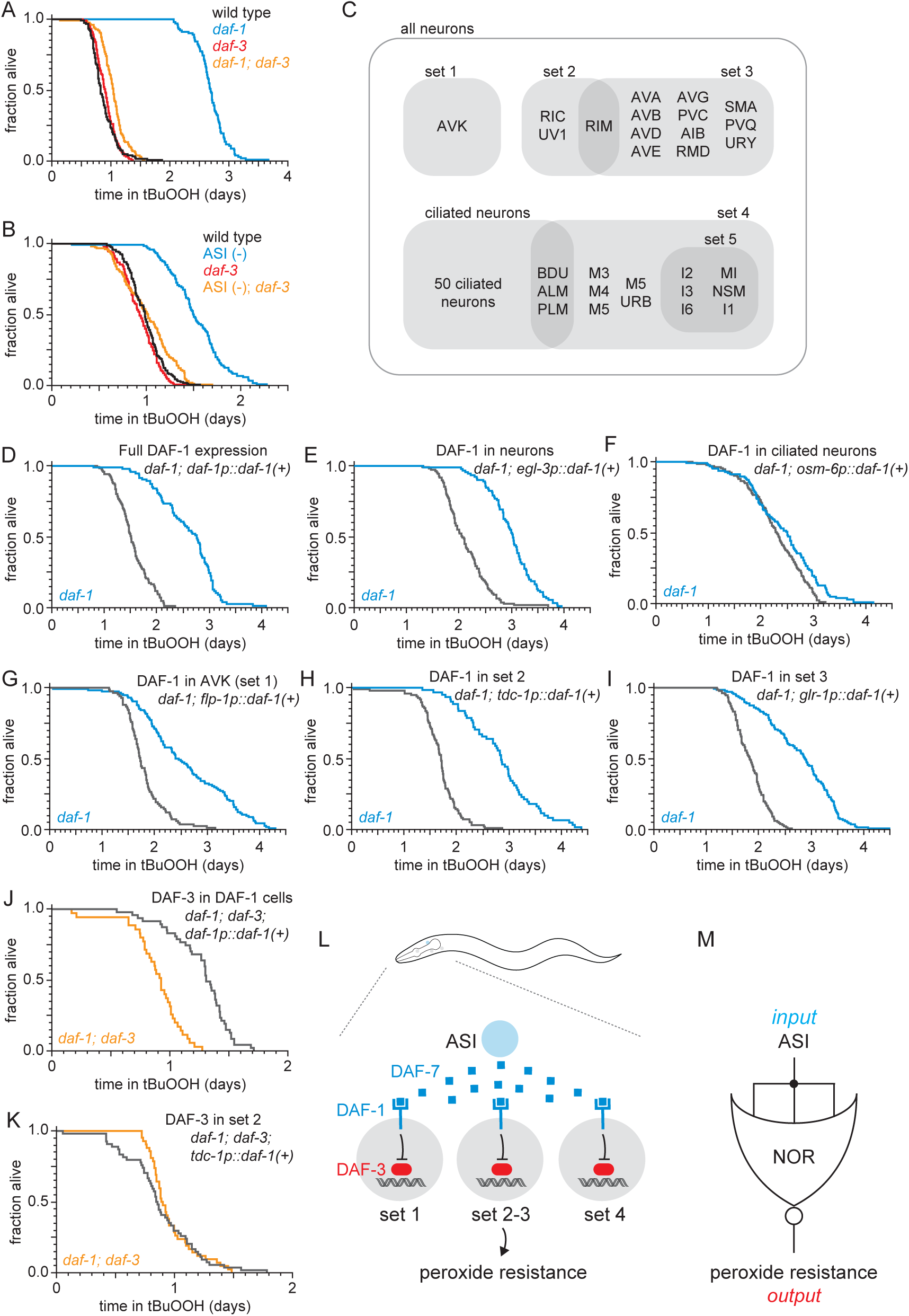
Interneurons must reach a consensus to increase peroxide resistance in response to DAF-7/TGFβ from ASI. (A-B) *daf-3(mgDf90)* almost completely abrogated the increased peroxide resistance of (A) *daf-1(m40)* and of (B) genetic ablation of ASI. (C) Diagram of the subsets of neurons where *daf-1(+)* or *daf-3(+)* was expressed in transgenic rescue experiments shown in panels (D-K) and Figure S3. The *tdc-1* promoter also drives expression in the sheath cells of the somatic gonad. (D-I) Peroxide resistance of transgenic nematodes expressing *daf-1(+)* in specific subsets of cells and *daf-1(m40)* controls. (J-K) Peroxide resistance of transgenic nematodes expressing *daf-3(+)* in specific subsets of cells and *daf-1(m40); daf-3(mgDf90)* controls. (L) ASI signals to three sets of interneurons to lower the nematode’s peroxide resistance. To increase peroxide resistance, all of these sets of neurons must independently activate the DAF-3/DAF-5 transcriptional complex. (M) ASI regulates the nematode’s peroxide resistance via NOR logic gate implemented by sets of interneurons acting in parallel to receive and invert the DAF-7 signal from ASI. See also Figure S3. Statistical analyses are in Table S3.

To determine which cells receive the DAF-7/TFGβ signal from the ASI neurons to regulate peroxide resistance, we restored *daf-1(+)* gene expression in specific subsets of neurons using cell-type specific promoters in *daf-1(m40)* mutants (Fig. 3C). DAF-1/TFGβ receptor is expressed broadly in the nervous system and in the distal-tip cells of the gonad (Gunther et al., 2000). Pan-neuronal expression of *daf-1(+)* with the *egl-3* promoter lowered peroxide resistance in *daf-1* mutants to the same extent as did expressing *daf-1(+)* with the endogenous *daf-1* promoter (Figs. 3D and 3E, and Table S3). Reconstituting *daf-1(+)* expression in all ciliated neurons (except BAG and FLP) using the *osm-6* promoter had a minimal effect on peroxide resistance (Fig. 3F and Table S3), indicating that *daf-1* functions in non-ciliated neurons. Expression of *daf-1(+)* in multiple sets of non-ciliated interneurons and pharyngeal neurons using the *flp-1*, *tdc-1*, *glr-1*, or *glr-8* promoters lowered peroxide resistance to a similar extent as pan-neuronal expression of *daf-1(+)* in *daf-1* mutants (Figs. 3G-I and S3D, and Table S3), while directed *daf-1(+)* expression in nine pharyngeal neurons using the *glr-7* promoter did not affect peroxide resistance (Fig. S3E and Table S3). The *flp-1* promoter is active only in the two AVK interneurons (Greer et al., 2008). In addition, the *flp-1*, *tdc-1*, *glr-1*, and *glr-8* promoters drive expression in non-overlapping cells, except for the expression overlap in the two RIM interneurons by the *tdc-1* and *glr-1* promoters (Greer et al., 2008) (Fig. 3C). We refer to the sets of neurons where *flp-1*, *tdc-1*, *glr-1*, and *glr-8* are expressed as “DAF-1-sufficiency sets”, because expression of *daf-1(+)* in any one of these sets of neurons is sufficient to lower the peroxide resistance of *daf-1* mutant nematodes. We conclude that DAF-1 functions redundantly in AVK interneurons and at least two other separate sets of neurons to lower the nematode’s peroxide resistance.

Because during signal transduction the DAF-1/TGFβ-receptor inhibits the DAF-3/coSMAD transcription factor, DAF-3 should be active only in neurons where signaling through DAF-1 is inactive. As a result, we expected that when DAF-1 is active only in one DAF-1-sufficiency set of neurons, DAF-3 should be active only in neurons outside that set, including the neurons of other non-overlapping DAF-1-sufficiency sets. This implied that to increase peroxide resistance DAF-3 must be active in all DAF-1-sufficiency sets of neurons. To test this prediction, we examined the effect on peroxide resistance of restoring *daf-3(+)* expression in just one of the DAF-1-sufficiency sets of neurons in *daf-1; daf-3* double mutants. Confirming our prediction, we found that restoring *daf-3(+)* expression with the *tdc-1* promoter was not sufficient to increase peroxide resistance in *daf-1; daf-3* double mutants (Figs. 3K and S3F, and Table S3). In contrast, the peroxide resistance of *daf-1; daf-3* double mutants increased upon restoring *daf-3(+)* expression in all four DAF-1-sufficiency sets of neurons with a *daf-1* promoter (Fig. 3J and Table S3). We conclude that the combination of the redundant action of DAF-1 in multiple sets of neurons and the repression of DAF-3 by DAF-1 in each of those neurons ensures that the nematode’s peroxide resistance stays low until all DAF-1-sufficiency sets of neurons de-repress DAF-3/coSMAD (Fig. 3L). The way these neurons determine the nematode’s peroxide resistance in response to DAF-7 signal levels is analogous to the way a logic NOR gate determines its output in response to its inputs. This circuit can be understood as an interneuron-consensus mechanism that ensures that target tissues induce a peroxide-protective response only if all DAF-1-sufficiency sets of neurons in the circuit fail to perceive the DAF-7 signal from ASI (Fig. 3M).

### ASI regulates the nematode’s peroxide resistance via a TGFβ-Insulin/IGF1 hormone relay

Previous studies have shown that distinct mechanisms act downstream of the DAF-3/coSMAD transcription factor to mediate the effects of DAF-7/TGFβ signaling on dauer-larva formation, fat storage, germline size, lifespan, and feeding (Dalfo et al., 2012; Greer et al., 2008; Shaw et al., 2007). Using a genetic approach, we asked whether DAF-7/TGFβ signaling acts via one or more of these mechanisms to regulate the nematode’s peroxide resistance (Fig. 4A).

**Figure 4.**
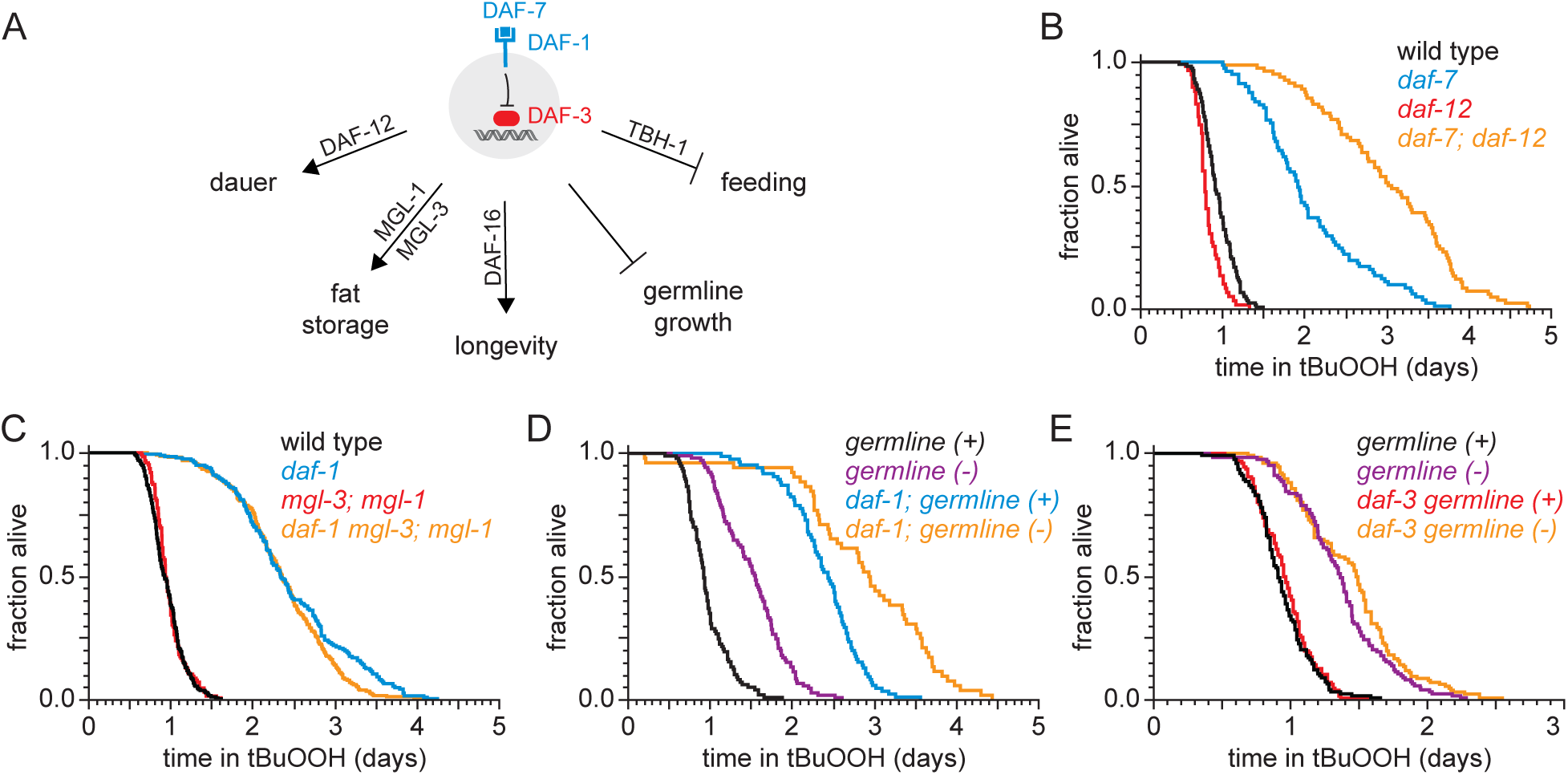
DAF-1/TGFβ-receptor signaling regulates peroxide resistance separately from its role in dauer formation, fat storage, and germline growth. (A) Different mechanisms operate downstream of DAF-3 to mediate the effects of DAF-7/TGFβ signaling on dauer-larva formation, fat storage, germline size, lifespan, and feeding. (B) *daf-12(rh61rh411)* did not suppress the increased peroxide resistance of *daf-7(e1372).* (C) *mgl-1(tm1811)* and *mgl-3(tm1766)* did not jointly suppress the increased peroxide resistance of *daf-1(m40).* (D) Genetic ablation of the germline and *daf-1(m40)* independently increased peroxide resistance. (E) *daf-3(mgDf90)* did not suppress the increased peroxide resistance of genetic ablation of the germline. See also Figure S4. Statistical analyses are in Table S4.

DAF-7 regulates dauer-larva formation via the nuclear hormone receptor DAF-12, which is the main switch driving the choice of reproductive growth or dauer arrest (Antebi et al., 2000). Loss of *daf-12* suppresses the constitutive dauer-formation phenotype of *daf-7* mutants during development (Thomas et al., 1993), but the *daf-12(rh61rh411)* null mutation did not suppress the increased peroxide resistance of *daf-7* mutant adults (Fig. 4B and Table S4). Therefore, DAF-7 regulates peroxide resistance and formation of peroxide-resistant dauer larvae via separate mechanisms.

The metabotropic glutamate receptors *mgl-1* and *mgl-3* are necessary for the increase in fat storage upon DAF-7-pathway inhibition (Greer et al., 2008). However, null mutations in either or both of these *mgl* genes did not affect peroxide resistance in *daf-1* mutants (Figs. 4C and S4, and Table S4). Thus, peroxide resistance and fat storage are also regulated via separate pathways downstream of DAF-1.

Germline size is reduced upon DAF-7-pathway inhibition (Dalfo et al., 2012). Genetic ablation of the germline increased the nematode’s peroxide resistance by 57% compared to wild-type animals (Fig. 4D and Table S4), consistent with previous studies (Steinbaugh et al., 2015). However, *daf-1(m40)* increased peroxide resistance in both germline-ablated and germline-non-ablated nematodes (Fig. 4D and Table S4). In addition, *daf-3(mgDf90)* did not affect peroxide resistance in germline-ablated nematodes (Fig. 4E and Table S4). Therefore, DAF-1 regulates peroxide resistance via a germline-independent mechanism.

Last, DAF-7-pathway signaling lowers lifespan by promoting insulin/IGF1 receptor signaling (Shaw et al., 2007). Previous studies have shown that transcription of at least 11 of the 40 insulin/IGF1 genes in the genome is repressed by the DAF-3/coSMAD in response to lower levels of DAF-7 and DAF1 signaling (Liu et al., 2004; Narasimhan et al., 2011; Shaw et al., 2007). We found that deletion of the DAF-3-repressed insulin/IGF1 genes *ins-1, ins-3, ins-4, ins-5, ins-6,* or *daf-28* caused increases in peroxide resistance ranging between 11% and 65% (Figs. 5A, S5A, and S5B, and Table S5), suggesting DAF-7 lowers peroxide resistance by promoting signaling by the insulin/IGF1 receptor, DAF-2. The *daf-2(e1370)* strong loss-of-function mutation increased peroxide resistance about three-fold (Fig. 5B and Table S5), consistent with previous findings (Tullet et al., 2008). Double mutants of *daf-1(m40)* and *daf-2(e1370)* had higher peroxide resistance than the respective single mutants (Fig. 5B and Table S5), suggesting that the DAF-1 TGFβ receptor and the DAF-2 insulin/IGF1 receptor regulate peroxide resistance via mechanisms that do not fully overlap. We considered the possibility that a DAF-2-dependent mechanism might mediate some of the effects of DAF-1 on peroxide resistance. If repressing the expression of insulin/IGF1 ligands of DAF-2 mediated part of the increased peroxide resistance of DAF-7-pathway inhibition, then one would expect the FOXO transcription factor DAF-16 to be necessary for those effects. DAF-16 is necessary for the increase in lifespan and most other phenotypes of mutants with reduced signaling by the DAF-2 insulin/IGF1 receptor (Kenyon et al., 1993; Lin et al., 1997; Ogg et al., 1997). We found that DAF-16 was also necessary for the increase in peroxide resistance of *daf-2(e1370)* mutants (Fig. 5C and Table S5) and for the increase in peroxide resistance of an *ins-4 ins-5 ins-6; daf-28* quadruple mutant (Fig. 5D and Table S5). The *daf-16(mu86)* null mutation decreased the peroxide resistance of *daf-7(e1372)* and *daf-1(m40)* mutants by nearly 50%, but caused only a small peroxide resistance reduction in wild-type nematodes (Figs. 5E and 5F, and Table S5). Therefore, regulation of peroxide resistance by the DAF-7/TGFβ signaling pathway is, in part, dependent on the DAF-16/FOXO transcription factor.

**Figure 5.**
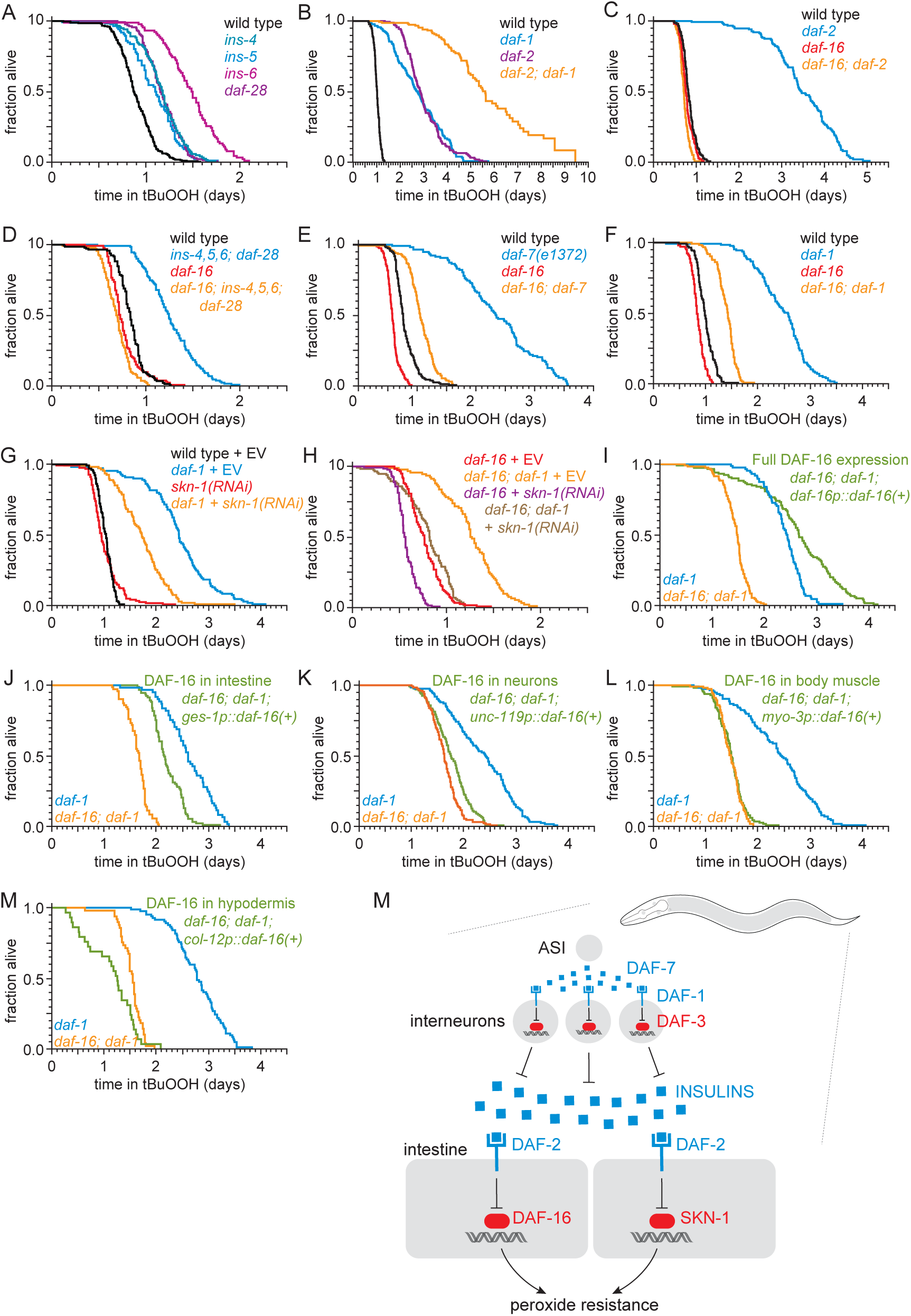
ASI regulates the nematode’s peroxide resistance via a TGFβ-Insulin/IGF1 hormone relay. (A) Deletions in *ins-4, ins-5, ins-6,* and *daf-28* insulin-coding genes increased peroxide resistance. (B) *daf-2(e1370)* and *daf-1(m40)* independently increased peroxide resistance. (C-D) *daf-16(mu86)* abrogated the increased peroxide resistance of (C) *daf-2(e1370)* and (D) an *ins-4 ins-5 ins-6; daf-28* quadruple mutant. (E-F) *daf-16(mu86)* suppressed part of the increased peroxide resistance of (E) *daf-7(e1372)* and (F) *daf-1(m40)*. (G) *skn-1(RNAi)* suppressed part of the increased peroxide resistance of *daf-1(m40)*. Control RNAi consisted of feeding the nematodes the same bacteria but with the empty vector (EV) plasmid pL4440 instead of a plasmid targeting *skn-1*. *(H) skn-1(RNAi)* lowered the peroxide resistance of *daf-16(mu86); daf-1(m40)*. (I-M) Peroxide resistance of transgenic nematodes expressing *daf-16(+)* in specific subsets of cells,*daf-16(mu86); daf-1(m40)* controls, and *daf-1(m40)* reference. (N) ASI sensory neurons make nematodes more sensitive to hydrogen peroxide via a multistep hormonal relay. DAF-7/TGFβ from ASI is received by interneurons. These interneurons act redundantly to relay this signal to target tissues by promoting transcription of insulin genes. These insulins activate the DAF-2 insulin/IGF1 receptor, leading to inhibition of DAF-16-dependent peroxide protection services by the intestine and neurons. SKN-1 acts independently of DAF-16 to promote peroxide resistance in response to reduced DAF-1 signaling. SKN-1 likely acts in the intestine, because *skn-1(+)* promotes peroxide resistance in *daf-2* mutants and induces oxidative-stress defenses in this tissue (An et al., 2005; Tullet et al., 2008). See also Figure S5. Statistical analyses are in Table S5.

We examined whether other transcription factors might act with DAF-16 to increase peroxide resistance in *daf-1* mutants. Like DAF-16, the NRF orthologue SKN-1 and the TFEB orthologue HLH-30 are activated in response to reduced DAF-2 signaling (Lin et al., 2018; Tullet et al., 2008). The peroxide resistance of *daf-1(m40) hlh-30(tm1978)* double mutants was identical to that of *daf-1* single mutants (Fig. S5C and Table S5). Knockdown of *skn-1* via RNA interference (RNAi) decreased the peroxide resistance of *daf-1(m40)* mutants by 30% but did not affect peroxide resistance in wild-type nematodes (Fig. 5G and Table S5). RNAi of *skn-1* also decreased the peroxide resistance of *daf-16; daf-1* double mutants, suggesting that DAF-16 and SKN-1 functioned in a non-overlapping manner to promote peroxide resistance in *daf-1(m40)* mutants (Fig. 5H and Table S5). We propose that repression of insulin/IGF1 gene expression by DAF-3/coSMAD leads to a reduction in signaling by the DAF-2/insulin/IGF1 receptor, which subsequently increases the nematode’s peroxide resistance via transcriptional activation by SKN-1/NRF and DAF-16/FOXO (Fig. 5N).

To identify which target tissues are important for increasing the nematode’s peroxide resistance via DAF-16 in response to reduced DAF-1 signaling, we determined the extent to which restoring *daf-16(+)* expression in specific tissues using tissue-specific promoters increased the peroxide resistance of *daf-16; daf-1* double mutants. As expected, peroxide resistance was increased when we restored *daf-16(+)* expression with the endogenous *daf-16* promoter (Fig. 5I and Table S5). Restoring *daf-16(+)* expression only in the intestine increased peroxide resistance, albeit to a lesser extent than did re-expressing *daf-16(+)* with the endogenous *daf-16* promoter (Fig. 5J and Table S5). Restoring *daf-16(+)* in neurons slightly increased peroxide resistance (Fig. 5K), while restoring *daf-16(+)* in body-wall muscles had no effect (Fig. 5L and Table S5). Restoring *daf-16(+)* expression in the hypodermis decreased peroxide resistance slightly (Fig. 5M and Table S5); however, it is difficult to interpret these results because these nematodes looked sickly (unlike *daf-1* and *daf-16* single and double mutants), consistent with reports that selectively expressing *daf-16(+)* in the hypodermis was toxic (Libina et al., 2003). Therefore, DAF-16/FOXO transcription in the intestine and neurons increases the nematode’s peroxide resistance when DAF-3/coSMAD is active due to reduced DAF-1 function (Fig. 5N).

### Food ingestion regulates the nematode’s peroxide resistance via DAF-3/coSMAD

Nematodes can be exposed directly to peroxides through food ingestion, and *daf-7* and *daf-1* mutants have been shown to exhibit mild feeding defects (Greer et al., 2008). Therefore, we considered the possibility that the increase in peroxide resistance of mutants with impaired DAF-7-pathway signaling was due to their reduced feeding. Previous studies have shown that the tyrosine decarboxylase TDC-1 and the tyramine β-hydroxylase TBH-1—biosynthetic enzymes for the neurotransmitters tyramine and octopamine, respectively—are each necessary for the feeding defect of *daf-1* mutants as *daf-1;tdc-1* and *daf-1;tbh-1* double mutants have normal feeding behaviors (Greer et al., 2008). Surprisingly, despite restoring normal feeding to *daf-1* mutants, *tbh-1* and *tdc-1* null mutations did not suppress the increased peroxide resistance of *daf-1* mutants (Figs. 6A and S6A, and Table S6). In fact, both mutations further increased peroxide resistance in a *daf-1* mutant background. Because mutations that restored normal feeding to *daf-1* mutants increased the peroxide resistance of *daf-1* mutants, these findings suggested that the reduced feeding exhibited by *daf-1* mutants in fact reduces the magnitude of their increased peroxide resistance.

To investigate whether feeding has a direct effect on peroxide resistance, we first determined whether a wild-type nematode’s feeding history (before peroxide exposure) might affect its subsequent peroxide resistance. We transferred nematodes to plates with different concentrations of *E. coli* for 24 hours prior to the start of the peroxide resistance assay and found that the *E. coli* concentration before the assay had a dose-dependent effect on peroxide resistance (Fig. 6B and Table S6). Animals grown on higher concentrations of *E. coli* had higher peroxide resistance. Strikingly, nematodes grown without *E. coli* for two days before the assay showed a six-fold decrease in peroxide resistance (Figs. 6C and S6B, and Table S6), even though they had access to plentiful *E. coli* during the assay.

**Figure 6.**
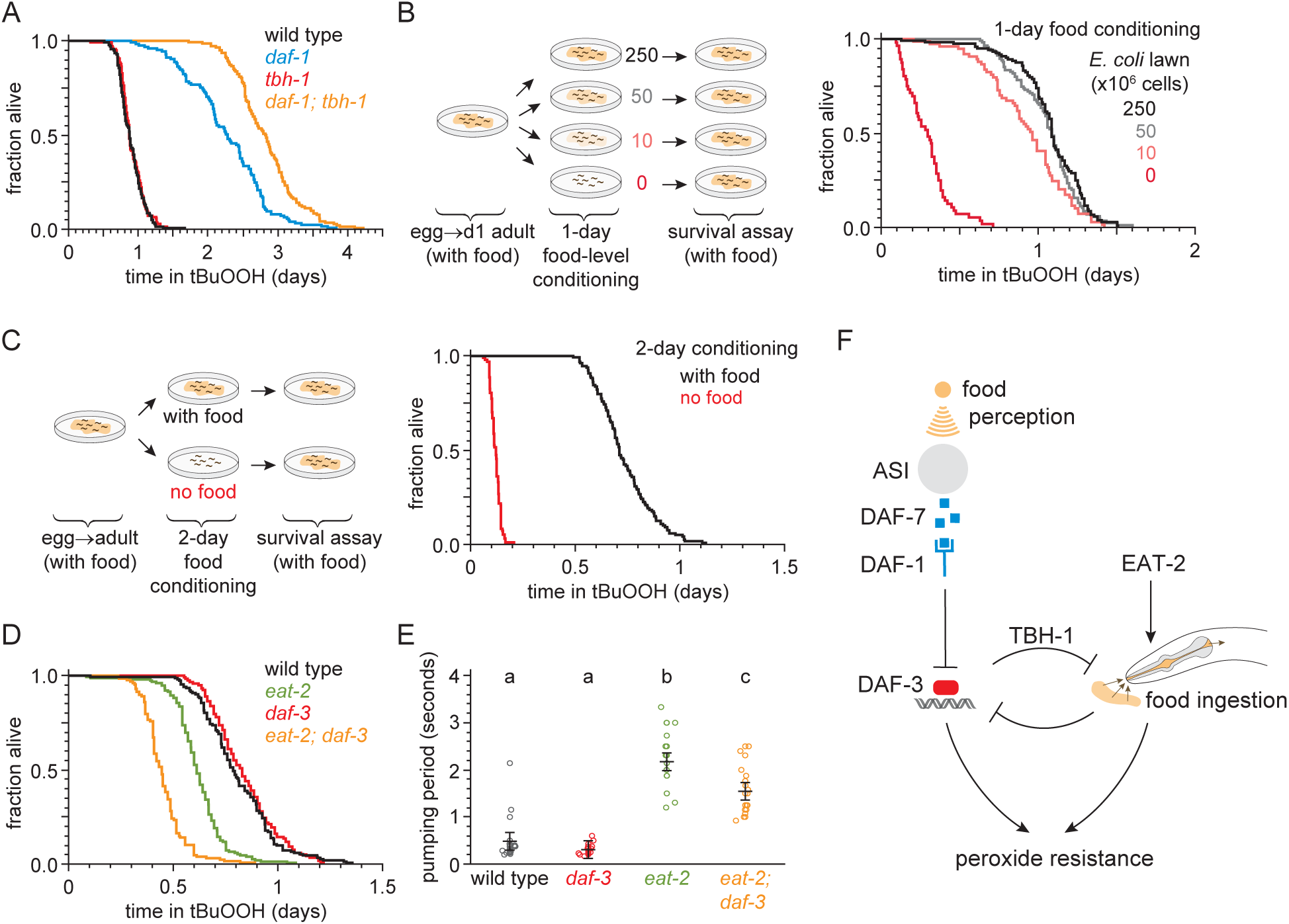
Food ingestion regulates the nematode’s peroxide resistance via DAF-3/coSMAD. (A) *tbh-1(ok1196)* increased the peroxide resistance of *daf-1(m40)*. (B-C) The *E. coli* level before the assay affected *C. elegans* peroxide resistance in a dose-dependent manner. (D) *eat-2(ad1116)* caused a more severe reduction in peroxide resistance in *daf-3(mgDf90)* than in wild type. (E) *eat-2(ad1116)* caused a less severe reduction in feeding in *daf-3(mgDf90)* than in wild type. Lines mark the mean pumping period and its 95% confidence interval. Genotypes labeled with different letters exhibited significant differences in pumping period (*P* < 0.0001, Turkey HSD test) otherwise (*P* > 0.05). (F) DAF-3 and feeding increase peroxide resistance but attenuate each other’s effects. Feeding inhibits DAF-3; this attenuates the reduction in peroxide resistance caused by reduced feeding. DAF-3 inhibits feeding via TBH-1; this attenuates the increase in peroxide resistance of *daf-1* mutants. Sensory perception of *E. coli* induces DAF-7 expression in (Chang et al., 2006; Gallagher et al., 2013) in a concentration-dependent manner (Entchev et al., 2015; Ren et al., 1996) leading to DAF-3 repression by the DAF-7 receptor DAF-1. Therefore, both ingestion and perception of *E. coli* inhibit DAF-3. See also Figure S6. Additional statistical analyses are in Table S6.

Next, we tested whether reduced ingestion of *E. coli* was sufficient to mimic the effects of pre-exposure to reduced *E. coli* levels. Mutants in the pharyngeal-muscle specific nicotinic acetylcholine receptor subunit *eat-2* ingest bacteria more slowly due to reduced pharyngeal pumping (feeding) (Avery, 1993; Raizen et al., 1995). The *eat-2(ad1116)* loss-of-function mutation, which causes a strong feeding defect (Fig. 6E), decreased peroxide resistance by 25% relative to wild-type animals (Fig. 6D and Table S6). Therefore, impaired feeding leads to decreased peroxide resistance.

Finally, we asked whether feeding and DAF-7 signaling regulate peroxide resistance jointly, or independently. Unlike *eat-2* mutants, *daf-3* null single mutants did not decrease peroxide resistance compared with wild-type animals (Figs. 3A, 3B, and 6D). However, the *eat-2(ad1116)* mutation caused a larger decrease in peroxide resistance in *daf-3* mutants than in wild-type nematodes (Fig. 6D and Table S6), suggesting that *daf-3(+)* promotes peroxide resistance in *eat-2* mutants. This effect was not due to an enhancement of the feeding defect of *eat-2* mutants by the *daf-3* mutation, because *eat-2; daf-3* double mutants fed slightly more (not less) than *eat-2* single mutants (Fig. 6E). We propose that DAF-3 is activated in response to reduced feeding, leading to an increase in peroxide resistance. DAF-3 acts as an adaptive mechanism that partially offsets the detrimental effect of reduced feeding on peroxide resistance.

Taken together, these findings imply that both feeding on bacteria and DAF-3/coSMAD signaling increase peroxide resistance, but that they attenuate each other’s effects (Fig. 6F). This cross-inhibition might enable nematodes to switch between DAF-3-dependent and DAF-3-independent mechanisms of peroxide resistance in response to changes in food ingestion and DAF-7 signal levels.

### DAF-7/TGFβ signals that hydrogen-peroxide protection will be provided by catalases from *E. coli* and not by catalases from *C. elegans*

Why does DAF-7 from ASI function to decrease the nematode’s peroxide resistance? ASI sensory neurons become active in response to perception of water-soluble signals from *E. coli* (Gallagher et al., 2013) and induce *daf-7* gene expression in a TAX-4-activity-dependent manner (Chang et al., 2006). As a result, the ASI neurons upregulate *daf-7* expression in response to *E. coli* (Gallagher et al., 2013) and lower *daf-7* gene expression in response to starvation and low *E. coli* concentrations (Entchev et al., 2015; Ren et al., 1996). Lowering DAF-7 levels when *E. coli* is scarce may enable nematodes to prepare for a future of reduced feeding by attenuating the expected reduction in peroxide resistance caused by reduced feeding. But increasing DAF-7 levels when *E. coli* is abundant may render nematodes more vulnerable to peroxide. We reasoned that perhaps *C. elegans* decreases its own peroxide self-defenses via DAF-7 signaling from the ASI neurons when *E. coli* is abundant because *C. elegans* expects to be safe from peroxide attack in that setting.

To test that hypothesis, we first asked whether *E. coli* can protect nematodes from the lethal effects of peroxides. This required that we re-examine the conditions of the peroxide resistance assays, which were conducted using a lipid hydroperoxide (tert-butyl hydroperoxide, tBuOOH) widely used in *C. elegans* studies due to its stability (An et al., 2005). When we used hydrogen peroxide instead of tBuOOH, we could not kill *C. elegans* even with concentrations as high as 20 mM (Fig. 7A) which is well above the biologically plausible range of up to 3 mM hydrogen peroxide used by other bacteria to kill *C. elegans* (Bolm et al., 2004; Moy et al., 2004). This suggested that hydrogen peroxide, but not tBuOOH, was efficiently degraded by *E. coli*. This bacterium uses several scavenging systems to degrade hydrogen peroxide (Mishra and Imlay, 2012). The two *E. coli* catalases, KatG and KatE, are the predominant scavengers of hydrogen peroxide in the environment, and the peroxiredoxin, AhpCF, plays a minor role (Seaver and Imlay, 2001). *E. coli* JI377, a *KatG KatE AhpCF* triple null mutant strain which cannot scavenge any hydrogen peroxide from the environment (Seaver and Imlay, 2001), did not protect *C. elegans* from 1 mM hydrogen peroxide killing (Fig. 7B), whereas the *E. coli* MG1655 parental wild-type strain was protective (Fig. 7B). We propose that *E. coli* protects *C. elegans* from hydrogen-peroxide killing because it expresses catalases that efficiently deplete hydrogen peroxide from the environment, creating a local environment where hydrogen peroxide is not a threat to *C. elegans*.

**Figure 7.**
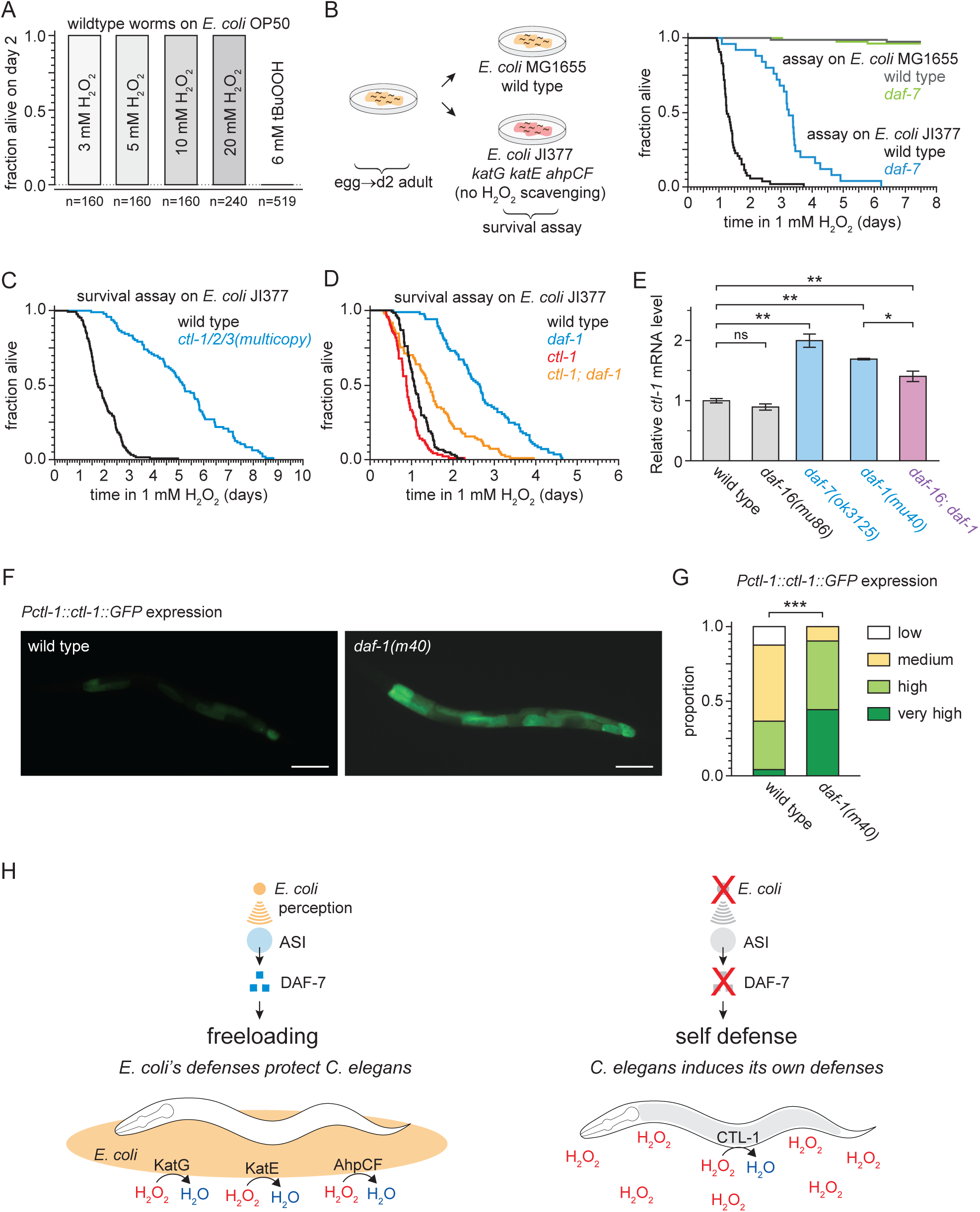
DAF-7/TGFβ signals that hydrogen-peroxide protection will be provided by catalases from *E. coli* and not by catalases from *C. elegans*. (A) *C. elegans* was sensitive to killing by tert-butyl hydroperoxide (tBuOOH), but not by hydrogen peroxide, in the presence of *E. coli* OP50. (B) Hydrogen peroxide resistance of wild type and *daf-7(ok3125*) *C. elegans* in assays with wild type and *Kat^−^ Ahp^−^ E. coli*. (C) Overexpression of the three endogenous catalases protects nematodes from hydrogen peroxide in assays with *Kat^−^ Ahp^−^ E. coli*. (D) The cytosolic catalase *ctl-1(ok1242)* mutation suppressed part of the increased hydrogen peroxide resistance of *daf-1(m40)* in assays with *Kat^−^ Ahp^−^ E. coli*. (E) DAF-7/TGFβ-pathway regulates *ctl-1* mRNA expression via DAF-16/FOXO, determined by quantitative RT-PCR. Data are represented as mean ± s.e.m of three independent biological replicates, each with three technical replicates. For comparisons of *ctl-1* mRNA expression between pairs of genotypes, ** indicates *P* < 0.001, * indicates *P* < 0.05, and “ns” indicates *P* > 0.05 (Turkey HSD test). (F) Representative pictures of the expression of the *chIs166[Pctl-1::ctl-1::gfp]* reporter in wild type animals (left picture; category: medium) or *daf-1(m40)* mutants (right picture; category: very high). Scale bar = 100 µm. (G) The expression of the promoter of *ctl-1* fused with GFP (*chIs166[Pctl-1::ctl-1::gfp]*) is higher in *daf-1(m40)* mutants (237 animals) than in wild type animals (145 animals), *** indicates *P* < 0.0001 (ordinal logistic regression). Scoring is described in Material and methods. See Figure S7C for representative pictures of each expression category. (H) DAF-7/TGFβ signaling enables *C. elegans* to decide whether to induce its own hydrogen-peroxide defenses or, instead, freeload on protection provided by molecularly orthologous hydrogen-peroxide defenses from *E. coli* See also Figure S7. Additional statistical analyses are in Table S7.

To determine whether DAF-7 regulates *C. elegans* hydrogen peroxide resistance, similar to its effects on tert-butyl hydroperoxide resistance, we examined resistance to hydrogen peroxide in *daf-7* mutants. In assays with the catalase mutant *E. coli* JI377 strain, we found that *daf-7(ok3125)* increased the nematode’s hydrogen peroxide resistance over two-fold relative to wild-type nematodes (Figure 7B and Table S7). ASI-ablation also increased hydrogen peroxide resistance in assays with *E. coli* JI377 (Fig. S7A and Table S7). We propose that in response to TAX-4-dependent sensory perception of *E. coli*, the ASI sensory neurons express DAF-7/TGFβ to instruct target tissues to downregulate their hydrogen peroxide defenses.

Last, we investigated the possibility that reducing DAF-7-pathway signaling protects *C. elegans* from hydrogen-peroxide killing via a hydrogen-peroxide defense mechanism orthologous to the one by which *E. coli* protects *C. elegans*. The *C. elegans* genome contains three catalase genes in tandem—two-newly duplicated cytosolic catalases, *ctl-1* and *ctl-3*, and a peroxisomal catalase, *ctl-2*—which are the nematode orthologues of the two *E. coli* catalases, *KatG* and *KatE* (Petriv and Rachubinski, 2004). We expected the *C. elegans* catalase genes to be upregulated in response to reduced DAF-7 signaling, because all three catalase genes have DAF-16 and SKN-1 binding sites in their promoters (An and Blackwell, 2003; Park et al., 2009; Petriv and Rachubinski, 2004), and their mRNA and protein expression increase in a DAF-16-dependent manner when DAF-2 signaling is reduced (Dong et al., 2007; McElwee et al., 2003; Murphy et al., 2003). To determine whether endogenous catalases could protect *C. elegans* from hydrogen peroxide when *E. coli* is not able to deplete hydrogen peroxide from the environment, we examined the effects of simultaneously increasing the dosage of all three catalase genes. We found that *ctl-1/2/3* overexpression, which increases catalase activity ten-fold (Doonan et al., 2008), more than doubled *C. elegans* hydrogen peroxide resistance in assays with *E. coli* JI377 (Fig. 7C and Table S7). To investigate whether one of the endogenous catalases might mediate the increased hydrogen peroxide resistance of nematodes with reduced DAF-7-pathway signaling, we constructed double mutants between *daf-1* and individual catalase genes. We found that the cytosolic catalase *ctl-1(ok1242)* null mutation abrogated much of the increase in hydrogen peroxide resistance of *daf-1(m40)* mutants in assays with *E. coli* JI377 (Fig. 7D and Table S7), but the peroxisomal catalase *ctl-2(ok1137)* null mutation did not (Fig. S7B and Table S7). Therefore, the increase in hydrogen peroxide resistance of *daf-1* mutants is mediated in part by the CTL-1 cytosolic catalase.

In line with this functional dependence, *ctl-1* mRNA levels were elevated up to two-fold in *daf-7(ok3125)* and *daf-1(m40)* mutant adults grown on *E. coli* OP50 (Fig. 7E). This upregulation was partially DAF-16-dependent, since the *daf-16(mu86)* mutation caused a small but statistically significant reduction in *ctl-1* mRNA expression in *daf-1(m40)* mutants but not in wild-type animals (Fig. 7E). The *ctl-1* gene product is expressed only in the intestine (Hamaguchi et al., 2019), and this expression was elevated in *daf-1(m40)* mutants (Figs. 7F and 7G). Taken together, these findings suggest that the DAF-7/TGFβ-pathway downregulates catalase gene expression in the intestine, partly via DAF-16. We propose that DAF-7/TGFβ signaling enables *C. elegans* to decide whether to induce its own hydrogen-peroxide degrading catalases or, instead, freeload on protection provided by molecularly orthologous catalases from *E. coli* (Fig. 7H).

## Discussion

Life forms throughout the evolutionary tree use hydrogen peroxide as an offensive weapon (Avery and Morgan, 1924; Imlay, 2018). Prevention and repair of the damage that hydrogen peroxide inflicts on macromolecules are critical for cellular health and survival (Chance et al., 1979). In this study, we found that in a simple animal, the nematode *C. elegans*, these protective responses are repressed in response to signals perceived by the nervous system. To our knowledge, the findings described here provide the first evidence of a multicellular organism modulating its defenses when it expects to freeload from the protection provided by molecularly orthologous defenses from individuals of a different species.

### Signals from sensory neurons regulate *C. elegans* hydrogen peroxide defenses

We show here that sensory neurons regulate how long *C. elegans* nematodes can survive in the presence of peroxides in the environment. Peroxide resistance was higher in nematodes with a global impairment in sensory perception (Fig. 1A). Using a systematic neuron-specific genetic-ablation approach, we identified ten classes of sensory neurons that influence the nematode’s peroxide resistance, including seven classes of neurons that normally decrease peroxide resistance and three classes of neurons that normally increase it (Fig. 1B). Why do so many neurons influence *C. elegans* peroxide resistance? One possibility is that these neurons respond to environments correlated with the threat of hydrogen peroxide.

Perception of water-soluble attractants by the amphid sensory neurons ASI, ASG, and ASK— neurons that when ablated caused some of the largest increases in peroxide resistance—helps *C. elegans* navigate towards bacteria (Bargmann and Horvitz, 1991a), its natural food source. We found that ingestion of *E. coli*, the nematode’s food in laboratory experiments, increases *C. elegans* peroxide resistance (Fig. 6D). In addition, *E. coli* expresses scavenging enzymes that degrade hydrogen peroxide in the nematode’s environment. These *E. coli* self-defense mechanisms create a public good (West et al., 2006), an environment safe from the threat of hydrogen peroxide, that benefits both *E. coli* and *C. elegans*. We propose that the control of organismic peroxide resistance by neurons that sense bacteria enables nematodes to turn down their peroxide self-defenses when they sense bacteria they deem protective. *C. elegans* freeloads off the hydrogen peroxide self-defense mechanisms from *E. coli* (Fig. 7H), because it uses a public good created by *E. coli*.

### The bacterial community influences the strategic choice between hydrogen peroxide self-defense and freeloading

We show here that *C. elegans* is safe from hydrogen peroxide attack when *E. coli* is abundant because hydrogen-peroxide degrading enzymes from *E. coli* protect *C. elegans*. *E. coli* degrades hydrogen peroxide in the environment primarily by expressing two catalases, KatG and KatE, as these enzymes account for over 95% of *E. coli’s* hydrogen-peroxide degrading capacity (Seaver and Imlay, 2001). Catalase-positive *E. coli* can protect catalase-deficient *E. coli* from hydrogen peroxide (Ma and Eaton, 1992). This facilitative relationship, where one species creates an environment that promotes the survival of another (Bronstein, 2009), also occurs across bacterial species in diverse environments: in dental plaque in the human mouth, *Actinomyces naeslundii* protects catalase-deficient *Streptococcus gordonii* by removing hydrogen peroxide (Jakubovics et al., 2008) and, in marine environments, catalase-positive bacteria protect the catalase-deficient cyanobacterium *Prochlorococcus*, the major photosynthetic organism in the open ocean (Zinser, 2018).

Unlike catalase-deficient bacteria receiving hydrogen-peroxide protection services from surrounding bacteria, *C. elegans* is not catalase deficient. In *C. elegans*, TAX-4-dependent sensory perception of *E. coli* stimulates the expression of DAF-7 in ASI (Chang et al., 2006; Entchev et al., 2015; Gallagher et al., 2013; Ren et al., 1996). We found that when DAF-7 signaling is reduced, target tissues induce defense mechanisms that protect *C. elegans* from hydrogen peroxide. These mechanisms are mediated in part by the DAF-16-dependent expression in the intestine of the cytosolic catalase CTL-1. We propose that the TGFβ-insulin/IGF1 signaling hormonal relay that begins with DAF-7 secretion from ASI enables this sensory neuron to communicate to target tissues that they do not need to induce CTL-1 and other hydrogen-peroxide protection services because *E. coli* in the surrounding environment likely provide molecularly orthologous services. Thus, this sensory circuit enables nematodes to choose between hydrogen-peroxide self-defense and freeloading strategies (Fig. 7H).

Hydrogen peroxide is a threat to *C. elegans*. In its natural habitat of rotting fruits and vegetation, *C. elegans* encounters a wide variety of bacterial taxa (Samuel et al., 2016), and this community includes bacteria in many genera known to degrade or produce hydrogen peroxide (Passardi et al., 2007). Hydrogen peroxide produced by a bacterium from the *C. elegans* microbiome, *Rhizobium huautlense*, causes DNA damage to the nematodes (Kniazeva and Ruvkun, 2019), and many bacteria—including *S. pyogenes*, *S. pneumoniae*, *S. oralis*, and *E. faecium*—kill *C. elegans* by producing hydrogen peroxide, often in concentrations exceeding the 1 mM concentration present in our assays (Bolm et al., 2004; Jansen et al., 2002; Moy et al., 2004). *C. elegans* may also encounter hydrogen peroxide derived from fruits, leaves, and stems, because plants produce hydrogen peroxide to attack their pathogens (Arakawa et al., 2014; Daudi et al., 2012 ; Mehdy, 1994). *C. elegans* also induces production of hydrogen peroxide to attack its pathogens, including *E. faecalis* (Chavez et al., 2007). In this complex and variable habitat, deciding whether to induce hydrogen-peroxide defenses is challenging. *C. elegans* cells manage this challenge by relinquishing control of their cellular hydrogen-peroxide defenses to a neuronal circuit in the nematode’s brain. This circuit might be able to integrate a wider variety of inputs than individual cells could, enabling a better assessment of the threat of hydrogen peroxide and precise regulation of hydrogen-peroxide protective defenses.

### Coordination of behavior, development, and physiology in response to the perceived threat of hydrogen peroxide

Assigning control of hydrogen peroxide cellular defenses to a sensory circuit in the brain could be beneficial because it enables *C. elegans* to avoid the energetic cost of unneeded protection. Tight control of hydrogen-peroxide defenses may also be necessary because a protective response might cause undesirable side effects. Nematodes overexpressing all three catalase genes exhibit a high level of mortality due to internal hatching of larvae, and this phenotype can be suppressed by joint overexpression of the superoxide dismutase SOD-1 (Doonan et al., 2008), an enzyme that produces hydrogen peroxide. While catalases can degrade large quantities of hydrogen peroxide, at low hydrogen peroxide concentrations these enzymes accumulate in the ferryl-radical intermediate of their catalytic cycle, which is a dangerous oxidizing agent (Imlay, 2013). Hydrogen peroxide is an important intracellular signaling molecule, and depletion of hydrogen peroxide by scavenging enzymes may interfere with signal transduction and affect cell behavior and differentiation (Veal et al., 2007).

Is the choice between hydrogen-peroxide self-defense and freeloading strategies regulated by DAF-7 limited to inducing hydrogen-peroxide protection services in target tissues? We favor an alternative possibility, that DAF-7 coordinates the induction of a broad phenotypic response to the perceived threat of hydrogen peroxide, because the phenotypic responses to lower DAF-7 signaling follow the expected desirable outcomes for animals that anticipate exposure to hydrogen peroxide: (i) re-routing development to form hydrogen-peroxide resistant dauer larva (Riddle and Albert, 1997); (ii) reducing proliferation of germline stem cells (Dalfo et al., 2012), to prevent hydrogen-peroxide induced damage to their DNA (Wyatt et al., 2017; Zong et al., 2014), (iii) reducing oocyte fertilization and egg-laying (McKnight et al., 2014; Trent, 1982), to increase the chances of progeny survival; (iv) reducing feeding (Greer et al., 2008), since many pathogenic bacteria produce hydrogen peroxide; (v) avoiding high oxygen concentrations (Chang et al., 2006), which are oxidizing; and (vi) increasing the nematode’s hydrogen peroxide resistance.

These diverse phenotypic responses might be triggered by different DAF-7 levels, reflecting the adaptive benefit of reducing the harm of hydrogen peroxide in each case. Perhaps for this reason, the DAF-7 signal is relayed via different circuits to target tissues mediating some of those responses. The DAF-1 receptor and the DAF-3/DAF-5 complex function in the somatic gonad to regulate germ-cell proliferation (Dalfo et al., 2012), and in RIM and RIC interneurons to regulate feeding, fat storage, egg laying, and dauer-larva formation (Greer et al., 2008). In contrast, to regulating hydrogen peroxide resistance, DAF-1 and DAF-3 function in at least three different sets of interneurons (Fig. 3L). One set includes RIM interneurons, and another comprises only the two AVK interneurons, which are not involved in regulating feeding, egg laying, and dauer-larva formation via DAF-1 signaling (Greer et al., 2008). The more complex role of interneuronal DAF-1 signaling in regulating hydrogen peroxide resistance suggests that *C. elegans* takes great care to avoid inducing hydrogen-peroxide protection services in target tissues unless DAF-7 levels are low.

### When do animals choose between freeloading and self-defense strategies?

Our studies provide a template for understanding how complex animals coordinate cellular hydrogen-peroxide defenses. We identify sensory neurons that respond to bacterial cues as important regulators of hydrogen-peroxide protection by *C. elegans* target tissues. Similar regulatory systems may exist in other animals. In mice, sensory neurons involved in pain perception respond to cues from *Staphylococcus aureus* by releasing neuropeptides that inhibit the activation of hydrogen-peroxide producing immune cells (Chiu et al., 2013), and some of the neuropeptides secreted by these sensory neurons, including galanin and calcitonin gene-related peptide, also induce hydrogen peroxide protection in target cells (Cui et al., 2010; Tullio et al., 2017). Assigning control of cellular defenses to dedicated sensory circuits may represent a general cellular-coordination tactic used by animals to regulate induction of self-defenses for hydrogen peroxide and perhaps other threats.

We show that the two ASI amphid sensory neurons use a multistep signal relay to control the extent to which target tissues protect *C. elegans* from hydrogen peroxide. The NOR circuit logic implemented by these sequential hormonal steps may enable ASI to control the induction of a sharp and specific peroxide-protective response in target tissues. In insects and mammals, TGFβ and insulin/IGF1 signaling components regulate cellular antioxidant defenses (Brunet et al., 2004; Clancy et al., 2001; Holzenberger et al., 2003; Kayanoki et al., 1994; Liu et al., 2012; Tatar et al., 2003), so it will be interesting to determine if a conserved hormonal relay controls hydrogen-peroxide defenses in all animals.

We delineate a neuronal circuit that processes sensory information to control the induction of hydrogen peroxide protection services by *C. elegans* target tissues. In fluctuating environments, we expect this circuit’s output (self-defense or freeloading) to provide an evolutionarily optimal strategy across its inputs (low or high *E. coli*) (Kussell and Leibler, 2005; Maynard Smith, 1982; Wolf et al., 2005). While a freeloading strategy may provide maximum fitness by inactivating self-defenses in environments where hydrogen peroxide is not a threat, this strategy need not provide maximum health or longevity to the organism. Consistent with this, in addition to lowering peroxide resistance in *C. elegans*, the ASI, ASG, and AWC amphid sensory neurons also shorten this organism’s lifespan in environments with no hydrogen peroxide (Alcedo and Kenyon, 2004), and DAF-7/TGFβ signaling from ASI also shortens *C. elegans* lifespan in those environments (Shaw et al., 2007). Thus, inducing latent self-defenses in environments where they are normally not induced can provide an approach to increase longevity in *C. elegans*. Because sensory perception and catalases also determine health and longevity in invertebrate and vertebrate animals (Apfeld and Kenyon, 1999; Libert et al., 2007; Murphy et al., 2007; Perez-Estrada et al., 2019; Riera et al., 2014), it is likely that sensory modulation presents a promising approach to induce latent defenses that could increase health and longevity in all animals.

## Materials and Methods

### *C. elegans* culture, strains, and transgenes

Wild-type *C. elegans* was Bristol N2. *C. elegans* were cultured on NGM agar plates seeded with *E. coli* OP50, unless noted otherwise. For a list of all bacterial and worm strains used in this study, see Table S8 and Table S9, respectively. Double and triple mutant worms were generated by standard genetic methods. For a list of PCR genotyping primers and enzymes, and phenotypes used for strain construction, see Table S10. The *Ptdc-1::daf-3(+)::GFP* (pKA533) and *Pdaf-1::daf-3(+)::GFP* (pKA534) plasmids (kindly provided by Kaveh Ashrafi) were injected at 30 ng/µl into *daf-1(m40) IV*; *daf-3(mgDf90) X* with 20 ng/µl *Pmyo-2::RFP* and 20 ng/µl *Punc-122::DsRed*, respectively, as co-injection markers.

### Survival assays

Automated survival assays were conducted using a *C. elegans* lifespan machine scanner cluster (Stroustrup et al., 2013). This platform enables the acquisition of survival curves with very high temporal resolution and large population sizes. All chemicals were obtained from Sigma. For hydrogen peroxide, tert-butyl hydroperoxide, sodium arsenite, paraquat, and dithiothreitol assays, the compound was added to molten agar immediately before pouring onto 50 mm NGM agar plates. Plates were dried (Stroustrup et al., 2013) and seeded with 100 µl of concentrated *E. coli* OP50 resuspended at an OD_600_ of 20 (Entchev et al., 2015). For RNAi experiments, the appropriate *E. coli* HT115 (DE3) strain was used instead. For hydrogen peroxide assays, *E. coli* MG1655 or JI377 were used instead (Seaver and Imlay, 2001). Nematodes were cultured at 20°C until the onset of adulthood, and then cultured at 25°C—to potentially enhance *daf-7* mutant phenotypes (Ren et al., 1996; Shaw et al., 2007)—in groups of up to 100, on plates with 10 μg/ml 5-fluoro-2-deoxyuridine (FUDR), to avoid vulval rupture (Leiser et al., 2016), prevent matricidal effects of *daf-7* pathway mutants (Shaw et al., 2007), and eliminate live progeny. As an alternative to FUDR, we inhibited formation of the eggshell of fertilized *C. elegans* embryos with RNAi of *egg-5* (Entchev et al., 2015), with identical results (Fig. S2A and S2C-D, and Table S2). For sodium arsenite, paraquat, and DTT assays, we adjusted the concentrations of these compounds to reduce the survival of wild-type nematodes about as much as in the peroxide survival assays. For experiments with *daf-1; daf-2* double mutants, which only develop as dauers at 20°C, all strains were grown at 15°C instead of 20°C until the onset of adulthood. For food-conditioning experiments, *E. coli* OP50 was resuspended in S Basal containing streptomycin (50 μg/ml) and seeded onto plates supplemented with both streptomycin and carbenicillin, each at 50 μg/ml, as described (Entchev et al., 2015). For *daf-1*, *daf-3*, and *daf-16* transgenic-rescue experiments, we picked only nematodes exhibiting bright expression of the respective GFP-fusion proteins. Day 2 adults were transferred to lifespan machine assay plates. A typical experiment consisted of up to four genotypes or conditions, with 4 assay plates of each genotype or condition, each assay plate containing a maximum of 40 nematodes, and 16 assay plates housed in the same scanner. All experiments were repeated at least once, yielding the same results. Scanner temperature was calibrated to 25°C with a thermocouple (ThermoWorks USB-REF) on the bottom of an empty assay plate. Death times were automatically detected by the lifespan machine’s image-analysis pipeline, with manual curation of each death time through visual inspection of all collected image data (Stroustrup et al., 2013), without knowledge of genotype or experimental condition.

### RNA interference

*E. coli* HT115 (DE3) bacteria with plasmids expressing dsRNA targeting specific genes were obtained from the Ahringer and Vidal libraries (Kamath et al., 2001; Rual et al., 2004). Empty vector plasmid pL4440 was used as control. Bacterial cultures were grown in LB broth with 100 μg/ml ampicillin at 37°C, induced with 0.1 M isopropyl-thiogalactopyranoside (IPTG) at 37°C for 4 hours, concentrated to an OD_600_ of 20, and seeded onto NGM agar plates containing 50 μg/ml carbenicillin and 2 mM IPTG.

### Quantitative RT-PCR

Total RNA was extracted from day 2 animals that were transferred at the L4 stage onto NGM agar plates with 10 μg/ml FUDR seeded with *E. coli* OP50 and grown at 25°C. RNA extraction and cDNA preparation were performed as described (Amrit et al., 2019). Quantitative RT-PCRs were performed using the Biorad CFX Connect machine. PCR reactions were undertaken in 96-well optical reaction plates (Bio-Rad Hard Shell PCR Plates). A 20 µl PCR reaction was set up in each well using the SYBR PowerUp Green Master Mix (Applied Biosystems, USA) with 10ng of the converted cDNA and 0.3 M primers. For each gene at least three independent biological samples were tested, each with three technical replicates. Primers used in this study include TTCCATTTCAAGCCTGCTC (*ctl-1* Fwd), ATAGTCTGGATCCGAAGAGG (*ctl-1* Rev), GGATTTGGACATGCTCCTC (*rpl-32* Fwd) (Amrit et al., 2019), and GATTCCCTTGCGGCTCTT (*rpl-32* Rev) (Amrit et al., 2019).

### Microscopy

Transgenic animals expressing a Bxy-CTL-1::GFP fusion under the control of the *C. elegans ctl-1* promoter (Hamaguchi et al., 2019) were scored at the young-adult stage using a fluorescence dissection stereomicroscope (Zeiss Discovery V12) under 100x magnification, following a scheme previously used to score a *gcs-1p::GFP* reporter with a similar pattern of intestinal expression (Wang et al., 2010). Low: only anterior or posterior intestine with patches of GFP. Medium: anterior and posterior intestine with patches GFP, middle of the intestine with dim GFP. High: anterior and posterior intestine with non-patchy GFP expression, middle of the intestine with patchy or dim GFP. Very high: strong and non-patchy GFP expression throughout the intestine. Fluorescence imaging was conducted as previously described (Romero-Aristizabal et al., 2014) with an Axioskop 2 FS plus microscope (Zeiss) equipped with a D470/20x excitation filter, a 500dcxr dichroic mirror, and a HQ535/50m emission filter (all from Chroma), using a Plan-Apochromat 10X 0.45 NA 2 mm working distance objective lens (1063-139, Zeiss). Young adult worms were placed on petri plates with modified Nematode Growth Media (to minimize background fluorescence) containing 6 mM levamisole to immobilize the animals (Romero-Aristizabal et al., 2014). Images were acquired with a Cool SNAP HQ^2^ 14-bit camera (Photometrics) at 4x4 binning and 20 ms exposure. We performed background subtraction by removing the mode intensity value of the entire image from each pixel. This procedure removes the background due to the agar and the camera noise, since most pixels in our images were part of the background. All microscopy was performed at 22°C.

### Behavioral assays

Pharyngeal pumping was assayed for 30 seconds on day 2 adults at 25°C using a dissecting microscope under 100x magnification.

### Statistical analysis

All statistical analyses were performed in JMP Pro version 14 (SAS). Survival curves were calculated using the Kaplan-Meier method. We used the log-rank test to determine if the survival functions of two or more groups were equal. For pumping-period assays, we used the Tukey HSD post-hoc test to determine which pairs of groups in the sample differ. For intestinal GFP expression assays, we used ordinal logistic regression to determine if expression levels were equal between groups.

## Acknowledgements

We thank Jennifer Whangbo, Phyllis Strauss, Veronica Godoy, and Erin Cram for critical reading and detailed comments on our manuscript. Joy Alcedo, Kaveh Ashrafi, Ryan Baugh, Denise Ferkey, Takaaki Hirotsu, James Imlay, Koichi Hasegawa, Jane Hubbard, Charlotte Kelley, Dennis Kim, Junho Lee, Andres Maricq, Roger Pocock, Piali Sengupta, Young-Jai You, and Yun Zhang kindly provided strains and plasmids. We benefitted from discussions with members of Erin Cram’s lab, Veronica Godoy, Edward Geisinger, and Yunrong Chai. Some strains were provided by the CGC, which is funded by NIH Office of Research Infrastructure Programs (P40 OD010440), and the National BioResources Project, Japan. The research was supported by National Science Foundation CAREER grant 1750065 to J.A. and National Institutes of Health grant R01AG051659 to A.G.

## Author contributions

J.S., F.S., W.H. and J.A. conceived and designed experiments, and analyzed data. J.S., F.S., S.S., S.J., J.S., H.T., S.B., N.Mc., A.V., and W.S. constructed strains and performed experiments. F.A performed Q-PCR experiments. J.A and A.G. provided guidance. J.S., F.S., and J.A. interpreted results and wrote the manuscript with contributions from the other authors.

## Competing interests

The authors declare that no competing interests exist.

