## Supplementary Information for "*Caenorhabditis elegans* processes sensory information to choose between freeloading and self-defense strategies"

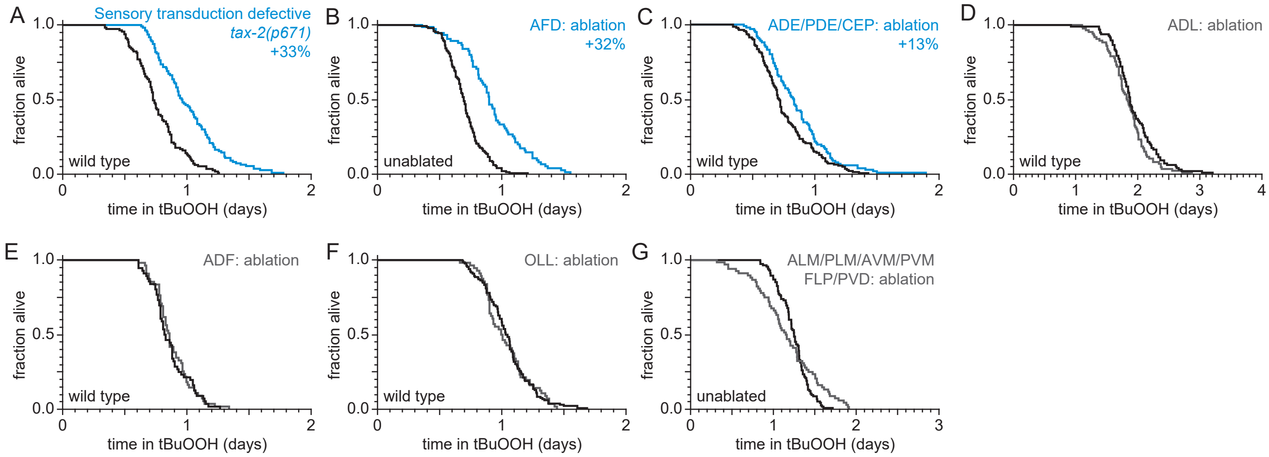


**Figure S1. Sensory neurons regulate peroxide resistance in *C. elegans*.**
(A-G) Peroxide resistance of nematodes with defects in sensory transduction, or with genetic ablation of specific sensory neurons. The fraction of nematodes remaining alive in the presence of 6 mM tert-butyl hydroperoxide (tBuOOH) is plotted against time. Interventions that increased or did not affect survival are denoted in blue and gray, respectively, and their effect on mean peroxide resistance is noted.
Statistical analyses are in Table S1.

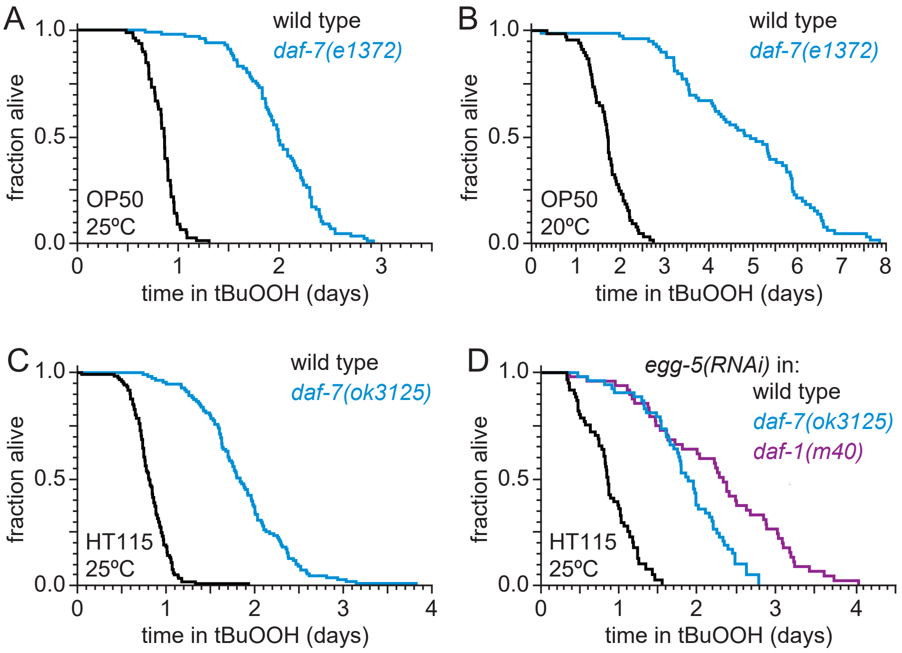


**Figure S2. DAF-7 regulates peroxide resistance robustly.**
(A-B) The *daf-7(e1372)* loss-of-function mutation increased peroxide resistance at (A) 25°C, and (B) 20°C.
(C) *daf-7(ok3125)* increased peroxide resistance when *C. elegans* was fed *E. coli* HT115(DE3) during development, adulthood, and the survival assay.
(D) *daf-7(ok3125)* and *daf-1(m40)* increase peroxide resistance when we inhibited formation of the eggshell of fertilized *C. elegans* embryos with RNAi of *egg*-5 and did not expose nematodes to FUDR at any point.
Statistical analyses are in Table S2.

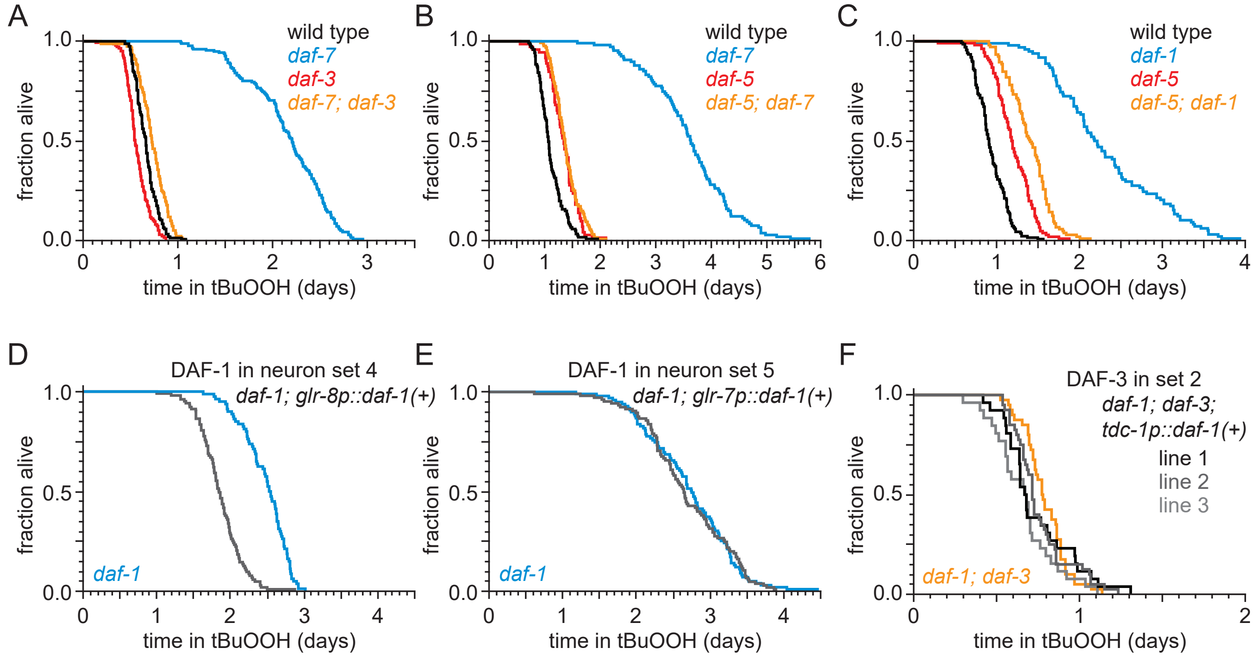


**Figure S3. Interneurons must reach a consensus to increase peroxide resistance in response to DAF-7/TGFβ from ASI.**

(A) *daf-3(e1376)* almost completely abrogated the increased peroxide resistance of *daf-7(ok3125)*.

(B) *daf-5(e1386)* abrogated the increased peroxide resistance of *daf-7(e1372)*.

(C) *daf-5(e1386)* abrogated part of the increased peroxide resistance of *daf-1(m40)*.

(D-E) Peroxide resistance of transgenic nematodes expressing *daf-1(+)* in specific subsets of cells and *daf-1(m40)* controls.

(F) Peroxide resistance of independent transgenic lines expressing *daf-3(+)* in *tdc-1*-expressing cells and *daf-1(m40); daf-3(mgDf90)* controls.

Statistical analyses are in Table S3.

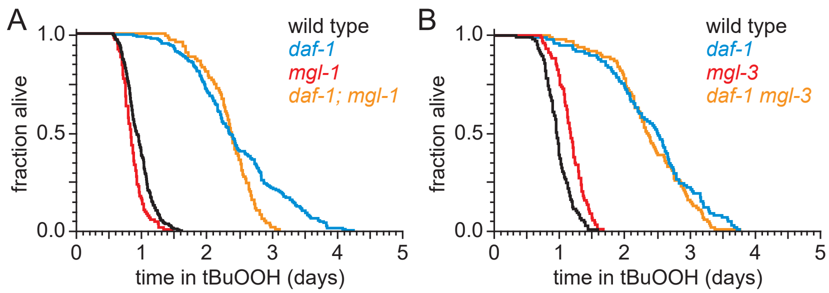


**Figure S4. DAF-1/TGFβ-receptor signaling regulates peroxide resistance separately from its role in fat storage.**

(A-B) The increased peroxide resistance of *daf-1(m40)* was not suppressed by (A) *mgl-1(tm1811)*, or (B) *mgl-3(tm1766).*

Statistical analyses are in Table S4.

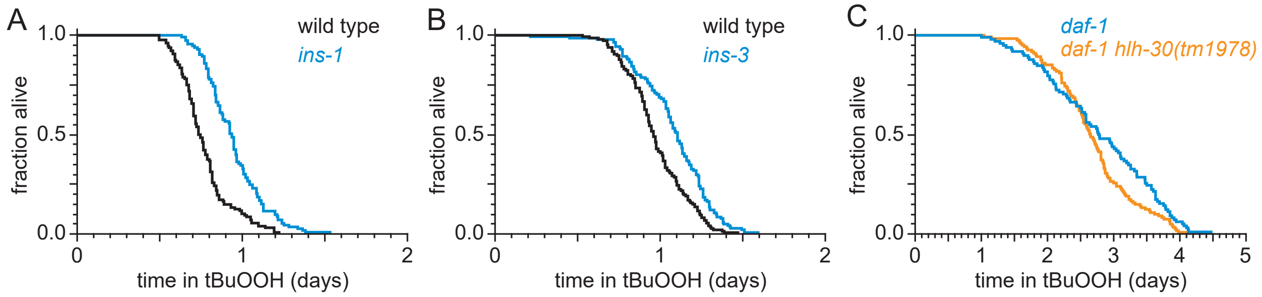


**Figure S5. ASI regulates the nematode's peroxide resistance via a TGFβ-Insulin/IGF1 hormone relay.**

(A-B) Peroxide resistance was increased by deletions in (A) *ins-1*, and (B) *ins-3* insulin-coding genes.
(C) *hlh-30(tm1978)* did not affect the increased peroxide resistance of *daf-1(m40)*.
Statistical analyses are in Table S5.

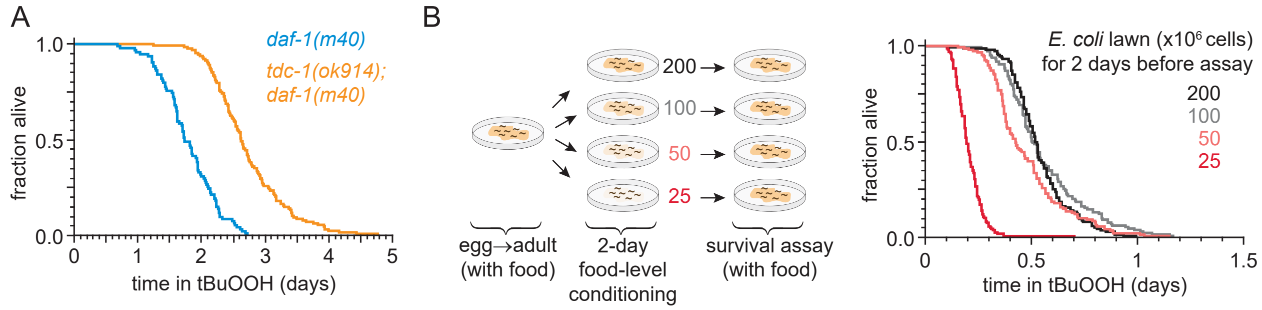


**Figure S6. DAF-3/coSMAD increases the nematode's peroxide resistance in response to reduced feeding.**

(A) *tdc-1(ok914)* increased the peroxide resistance of *daf-1(m40)*.

(B) The *E. coli* level for two days before the assay affected *C. elegans* peroxide resistance in a dose-dependent manner.

Statistical analyses are in Table S6.

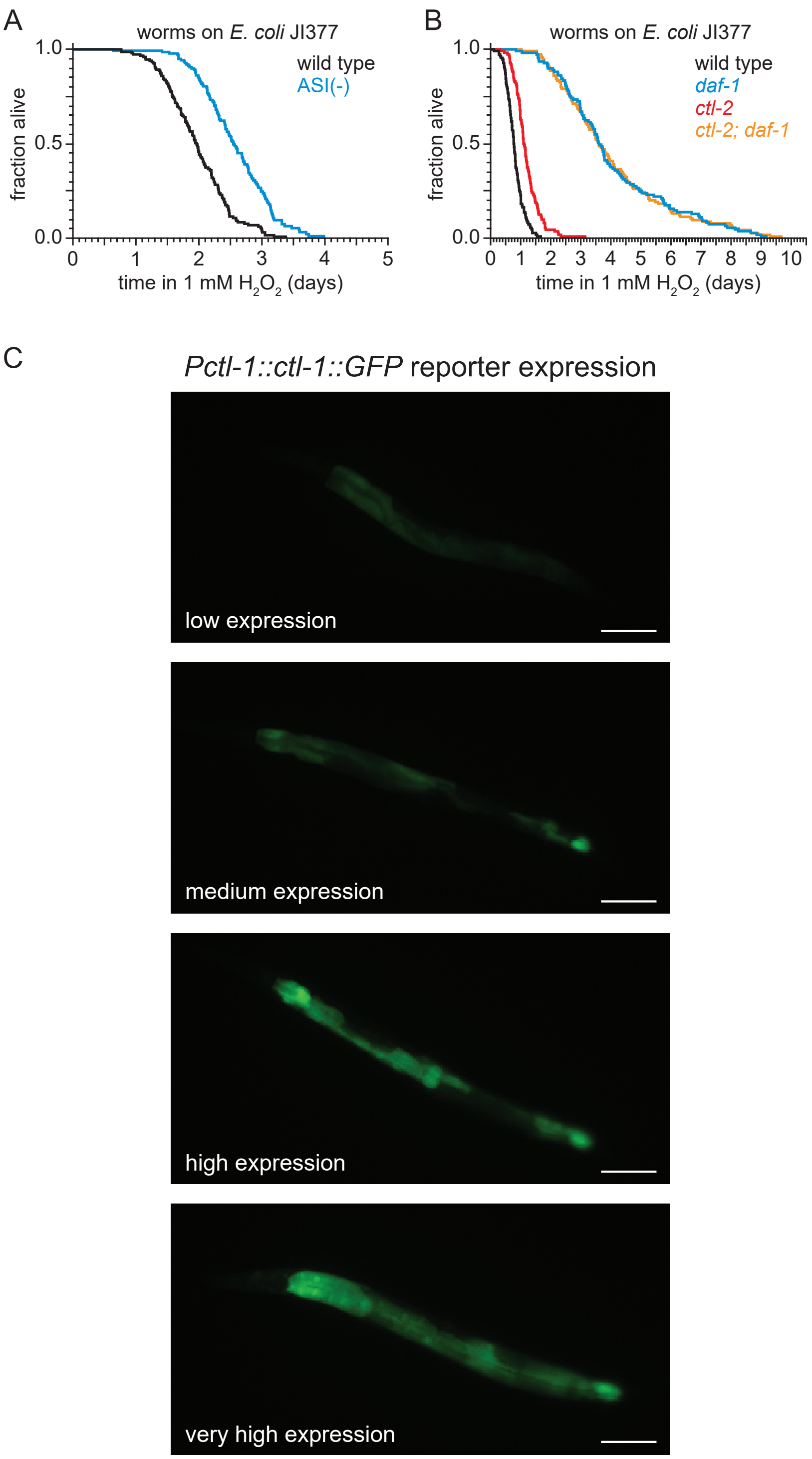


**Figure S7. DAF-7/TGFβ signals that hydrogen-peroxide protection will be provided by catalases from *E. coli* and not by catalases from *C. elegans*.**

(A) Genetic ablation of ASI sensory neurons increased peroxide resistance.

(B) The peroxisomal catalase *ctl-2(ok1137)* did not suppresses the increased peroxide resistance of *daf-1(m40)*.

(C) Representative pictures of the expression of the *chIs166[Pctl-1::ctl-1::gfp]* reporter. Scale bar = 100 µm.

Statistical analyses are in Table S7.

#### Table S1. Statistical analysis for Figures 1 and S1.

| Set | Genotype | Mean survival ± SEM (days) | Median survival (days) | 75th percentile (days) | N dead / initial N | % Mean survival change vs. control | *P* value (log-rank) vs. control | Figure |
|---|---|---|---|---|---|---|---|---|
| Cilium structure | | | | | | | | |
| | *osm-5(p813) X* | 1.06 ± 0.02 | 1.05 | 1.21 | 123 / 123 | 45% | < 0.0001 | 1 |
| | wild type | 0.73 ± 0.02 | 0.70 | 0.83 | 118 / 118 | | | |
| Sensory transduction | | | | | | | | |
| | *tax-4(p678) III* | 1.61 ± 0.03 | 1.64 | 1.79 | 138 / 138 | 124% | < 0.0001 | 1 |
| | wild type | 0.72 ± 0.02 | 0.68 | 0.83 | 111 / 111 | | | |
| | *tax-2(p671) I* | 1.01 ± 0.02 | 0.96 | 1.16 | 110 / 110 | 33% | < 0.0001 | S1A |
| | wild type | 0.76 ± 0.02 | 0.73 | 0.88 | 113 / 113 | | | |
| Neuronal genetic ablation | | | | | | | | |
| | ASG: ablation | 1.19 ± 0.02 | 1.21 | 1.32 | 138 / 149 | 48% | < 0.0001 | 1 |
| | wild type | 0.80 ± 0.02 | 0.75 | 0.90 | 127 / 143 | | | |
| | ASE: *che-1(ot75) I* | 0.91 ± 0.01 | 0.91 | 1.00 | 138 / 138 | 2% | > 0.05 | 1 |
| | wild type | 0.89 ± 0.01 | 0.89 | 0.98 | 144 / 144 | | | |
| | ASI: ablation | 1.41 ± 0.02 | 1.44 | 1.50 | 117 / 128 | 61% | < 0.0001 | 1 |
| | wild type | 0.88 ± 0.01 | 0.86 | 0.98 | 130 / 138 | | | |
| | ASK: ablation | 0.99 ± 0.02 | 0.96 | 1.20 | 129 / 141 | 13% | < 0.0001 | 1 |
| | wild type | 0.88 ± 0.02 | 0.87 | 1.00 | 134 / 144 | | | |
| | IL2: ablation | 1.46 ± 0.03 | 1.39 | 1.71 | 131 / 131 | 39% | < 0.0001 | 1 |
| | wild type | 1.06 ± 0.02 | 1.05 | 1.17 | 131 / 131 | | | |
| | ASJ: ablation | 1.16 ± 0.03 | 1.10 | 1.33 | 82 / 82 | -14% | 0.0016 | 1 |
| | unablated (no transgene) | 1.34 ± 0.04 | 1.35 | 1.63 | 93 / 93 | | | |
| | AFD: ablation | 1.29 ± 0.03 | 1.27 | 1.47 | 138 / 138 | 39% | < 0.0001 | 1 |
| | wild type | 0.92 ± 0.02 | 0.89 | 1.06 | 137 / 137 | | | |
| | AFD: ablation (transgene) | 1.39 ± 0.04 | 1.35 | 1.65 | 75 / 75 | 32% | < 0.0001 | S1B |
| | unablated (no transgene) | 1.05 ± 0.02 | 1.04 | 1.18 | 147 / 147 | | | |
| | AWA: ablation | 0.71 ± 0.02 | 0.71 | 0.83 | 72 / 72 | -16% | < 0.0001 | 1 |
| | unablated (no transgene) | 0.84 ± 0.02 | 0.86 | 0.95 | 80 / 80 | | | |
| | AWB: ablation | 0.78 ± 0.02 | 0.77 | 0.91 | 106 / 106 | -4% | > 0.05 | 1 |
| | wild type | 0.82 ± 0.02 | 0.81 | 0.94 | 114 / 114 | | | |
| | AWC: ablation | 0.89 ± 0.02 | 0.88 | 1.02 | 137 / 137 | 8% | 0.0037 | 1 |
| | wild type | 0.83 ± 0.01 | 0.83 | 0.92 | 139 / 139 | | | |
| | ASH: ablation | 0.89 ± 0.02 | 0.89 | 1.01 | 87 / 87 | 4% | > 0.05 | 1 |
| | wild type | 0.86 ± 0.02 | 0.83 | 0.95 | 128 / 128 | | | |
| | ADE/PDE/CEP: ablation | 0.85 ± 0.03 | 0.83 | 0.99 | 101 / 101 | 13% | 0.0065 | S1C |
| | wild type | 0.75 ± 0.02 | 0.71 | 0.88 | 140 / 140 | | | |
| | ADL: ablation | 1.83 ± 0.04 | 1.83 | 2.01 | 86 / 86 | -6% | > 0.05 | S1D |
| | wild type | 1.94 ± 0.04 | 1.88 | 2.13 | 97 / 97 | | | |
| | ADF: ablation | 0.88 ± 0.02 | 0.86 | 0.98 | 55 / 55 | 2% | > 0.05 | S1E |
| | wild type | 0.86 ± 0.02 | 0.82 | 0.94 | 56 / 56 | | | |
| | OLL: ablation | 1.04 ± 0.03 | 1.00 | 1.14 | 56 / 56 | -1% | > 0.05 | S1F |
| | unablated (no transgene) | 1.05 ± 0.02 | 1.04 | 1.15 | 132 / 132 | | | |
| | ALM/PLM/AVM/PVM/FLP/PVD: ablation (transgene) | 1.17 ± 0.05 | 1.16 | 1.45 | 68 / 68 | -6% | > 0.05 | S1G |
| | unablated (no transgene) | 1.24 ± 0.02 | 1.25 | 1.36 | 115 / 115 | | | |
| | URX/AQR/PQR: ablation | 0.83 ± 0.02 | 0.79 | 0.98 | 145 / 145 | -15% | < 0.0001 | 1 |
| | wild type | 0.98 ± 0.02 | 0.97 | 1.12 | 142 / 142 | | | |

### Table S2. Statistical analysis for Figures 2 and S2.

| Set | Genotype | Mean survival ± SEM (days) | Median survival (days) | 75th percentile (days) | N dead / initial N | % Mean survival change vs. wild type | Group | *P* value (log-rank) vs. wild type | *P* value (log-rank) vs. group b | *P* value (log-rank) vs. group c | Figure |
|---|---|---|---|---|---|---|---|---|---|---|---|
| 6 mM tBuOOH, 25°C, OP50 | | | | | | | | | | | |
| | wild type | 0.92 ± 0.02 | 0.90 | 1.02 | 98 / 98 | | a | | | | |
| | *daf-7(ok3125) III* (no *qdEx37* transgene) | 1.88 ± 0.04 | 1.94 | 2.10 | 83 / 83 | 104% | b | < 0.0001 | | | 2B |
| | *daf-7(ok3125) III; qdEx37[Pdaf-7::daf-7(+), Pges-1::GFP* | 0.97 ± 0.02 | 0.95 | 1.06 | 117 / 117 | 5% | c | > 0.05 | < 0.0001 | < 0.0001 | |
| | wild type | 0.97 ± 0.02 | 0.94 | 1.10 | 109 / 109 | | a | | | | |
| | *daf-7(ok3125) III* (no *qdEx44* transgene) | 2.35 ± 0.05 | 2.41 | 2.66 | 89 / 89 | 142% | b | < 0.0001 | | | |
| | *daf-7(ok3125) III; qdEx44[Pstr-3::daf-7(+), Pges-1::GFP]* | 1.05 ± 0.03 | 1.05 | 1.17 | 80 / 80 | 9% | c | 0.0031 | < 0.0001 | < 0.0001 | 2C,D |
| | *daf-7(ok3125) III* (no q*dEx40* transgene) | 2.21 ± 0.07 | 2.32 | 2.50 | 44 / 44 | 128% | d | < 0.0001 | | | |
| | *daf-7(ok3125) III; qdEx40[Ptrx-1::daf-7(+), Pges-1::GFP]* | 0.99 ± 0.02 | 0.98 | 1.11 | 70 / 70 | 2% | e | > 0.05 | < 0.0001 | < 0.0001 | |
| | wild type | 0.84 ± 0.02 | 0.86 | 0.92 | 78 / 81 | | a | | | | S2A |
| | *daf-7(e1372) III* | 1.99 ± 0.04 | 1.99 | 2.29 | 94 / 101 | 137% | b | < 0.0001 | | | |
| 6 mM tBuOOH, 20°C, OP50 | | | | | | | | | | | |
| | wild type | 1.71 ± 0.06 | 1.73 | 1.98 | 67 / 68 | | a | | | | S2B |
| | *daf-7(e1372) III* | 4.81 ± 0.17 | 4.95 | 5.90 | 73 / 79 | 181% | b | < 0.0001 | | | |
| 6 mM tBuOOH, 25°C, HT115 | | | | | | | | | | | |
| | wild type | 0.82 ± 0.02 | 0.81 | 0.94 | 121 / 121 | | a | | | | S3B |
| | *daf-7(e1372) III* | 1.87 ± 0.05 | 1.84 | 2.17 | 111 / 111 | 127% | b | < 0.0001 | | | |
| 6 mM tBuOOH, 25°C, HT115 *egg-5(RNAi)* , no FUDR | | | | | | | | | | | |
| | wild type | 0.86 ± 0.04 | 0.85 | 1.13 | 55 / 61 | | a | | | | |
| | *daf-7(ok3125) III* | 1.85 ± 0.08 | 1.87 | 2.23 | 48 / 53 | 113% | b | < 0.0001 | | | S4B |
| | *daf-1(m40) IV* | 2.25 ± 0.13 | 2.31 | 3.00 | 46 / 52 | 160% | c | < 0.0001 | | | |
| 5 mM arsenite, 25°C, OP50 | | | | | | | | | | | |
| | wild type | 1.27 ± 0.02 | 1.26 | 1.37 | 117 / 117 | | a | | | | 2E |
| | *daf-7(ok3125) III* | 1.22 ± 0.03 | 1.23 | 1.42 | 111 / 111 | -4% | b | > 0.05 | | | |
| 75 mM paraquat, 25°C, OP50 | | | | | | | | | | | |
| | wild type | 1.97 ± 0.05 | 2.00 | 2.37 | 90 / 90 | | a | | | | 2F |
| | *daf-7(ok3125) III* | 2.11 ± 0.07 | 2.14 | 2.57 | 93 / 93 | 7% | b | > 0.05 | | | |
| 25 mM dithiothreitol (DTT), 25°C, OP50 | | | | | | | | | | | |
| | wild type | 0.37 ± 0.01 | 0.36 | 0.42 | 111 / 116 | | a | | | | 2G |
| | *daf-7(ok3125) III* | 0.40 ± 0.01 | 0.39 | 0.48 | 110 / 110 | 7% | b | > 0.05 | | | |

### Table S3 . Statistical analysis for Figures 3 and S3.

| Set | Genotype | Mean survival ± SEM (days) | Median survival (days) | 75th percentile (days) | N dead / initial N | Group | % Mean survival change vs. group a | *P* value (log-rank) vs. group a | *P* value (log-rank) vs. group b | *P* value (log-rank) vs. group c | Figure |
|---|---|---|---|---|---|---|---|---|---|---|---|
| Pathway analysis | | | | | | | | | | | |
| | wild type | 0.88 ± 0.02 | 0.82 | 0.99 | 122 / 122 | a | | | | | 3A |
| | *daf-1(m40) IV* | 2.69 ± 0.03 | 2.69 | 2.84 | 102 / 102 | b | 207% | < 0.0001 | | | |
| | *daf-3(mgDf90) X* | 0.91 ± 0.02 | 0.90 | 1.04 | 130 / 130 | c | 3% | > 0.05 | < 0.0001 | | |
| | *daf-1(m40) IV; daf-3(mgDf90) X* | 1.04 ± 0.02 | 1.04 | 1.15 | 127 / 127 | d | 19% | < 0.0001 | < 0.0001 | < 0.0001 | |
| | wild type | 0.66 ± 0.01 | 0.66 | 0.73 | 156 / 163 | a | | | | | S3A |
| | *daf-7(ok3125) III* | 2.15 ± 0.03 | 2.21 | 2.49 | 154 / 176 | b | 223% | < 0.0001 | | | |
| | *daf-3(e1376) X* | 0.57 ± 0.01 | 0.55 | 0.64 | 139 / 143 | c | -14% | < 0.0001 | < 0.0001 | | |
| | *daf-7(ok3125) III; daf-3(e1376) X* | 0.73 ± 0.01 | 0.73 | 0.84 | 151 / 157 | d | 10% | < 0.0001 | < 0.0001 | < 0.0001 | |
| | wild type | 1.11 ± 0.02 | 1.07 | 1.25 | 121 / 121 | a | | | | | S3B |
| | *daf-7(e1372) III* | 3.63 ± 0.08 | 3.65 | 4.17 | 107 / 107 | b | 226% | < 0.0001 | | | |
| | *daf-5(e1386) II* | 1.34 ± 0.03 | 1.35 | 1.51 | 72 / 72 | c | 21% | < 0.0001 | < 0.0001 | | |
| | *daf-5(e1386) II; daf-7(e1372) III* | 1.39 ± 0.02 | 1.38 | 1.54 | 128 / 128 | d | 25% | < 0.0001 | < 0.0001 | > 0.05 | |
| | wild type | 0.93 ± 0.02 | 0.91 | 1.08 | 137 / 137 | a | | | | | S3C |
| | *daf-1(m40) IV* | 2.32 ± 0.07 | 2.20 | 2.77 | 98 / 98 | b | 149% | < 0.0001 | | | |
| | *daf-5(e1386) II* | 1.20 ± 0.02 | 1.41 | 1.59 | 106 / 106 | c | 29% | < 0.0001 | < 0.0001 | | |
| | *daf-5(e1386) II; daf-1(m40) IV* | 1.40 ± 0.02 | 1.19 | 1.38 | 105 / 105 | d | 50% | < 0.0001 | < 0.0001 | < 0.0001 | |
| | wild type | 0.99 ± 0.01 | 0.99 | 1.10 | 161 / 161 | a | | | | | 3B |
| | ASI ablation | 1.52 ± 0.03 | 1.50 | 1.71 | 120 / 120 | b | 54% | < 0.0001 | | | |
| | *daf-3(mgDf90) X* | 0.93 ± 0.02 | 0.93 | 1.07 | 150 / 150 | c | -6% | 0.03 | < 0.0001 | | |
| | ASI ablation; *daf-3(mgDf90) X* | 1.00 ± 0.02 | 1.02 | 1.20 | 156 / 156 | d | 2% | 0.0045 | < 0.0001 | < 0.0001 | |
| Full DAF-1 expression | | | | | | | | | | | |
| | *daf-1(m40) IV* (no *ftEx98* transgene) | 2.58 ± 0.06 | 2.76 | 3.00 | 92 / 94 | a | | | | | 3D |
| | *daf-1(m40) IV; ftEx98[Pdaf-1::daf-1-gfp Podr-1::dsRED]* | 1.54 ± 0.03 | 1.51 | 1.76 | 115 / 121 | b | -40% | < 0.0001 | | | |
| DAF-1 in neurons | | | | | | | | | | | |
| | *daf-1(m40) IV (no ftEx69 transgene)* | 3.00 ± 0.05 | 3.03 | 3.29 | 91 / 91 | a | | | | | 3E |
| | *daf-1(m40); ftEx69[Pegl-3::daf-1::GFP + Podr-1::dsRED]* | 2.13 ± 0.04 | 2.06 | 2.38 | 109 / 109 | b | -29% | < 0.0001 | | | |
| DAF-1 in ciliated neurons | | | | | | | | | | | |
| | *daf-1(m40) IV* (no *ftEx83* transgene) | 2.41 ± 0.06 | 2.51 | 2.89 | 114 / 126 | a | | | | | 3F |
| | *daf-1(m40ts) IV; ftEx83[Posm-6::daf-1-gfp + Podr-1::dsRED]* | 2.27 ± 0.05 | 2.30 | 2.69 | 128 / 133 | b | -6% | 0.0048 | | | |
| DAF-1 in AVK (set 1) | | | | | | | | | | | |
| | *daf-1(m40) IV* (no *ftEx166* transgene) | 2.58 ± 0.08 | 2.43 | 3.33 | 114 / 114 | a | | | | | 3G |
| | *daf-1(m40) IV; ftEx166[Pflp-1::daf-1::gfp + Podr-1::dsRED]* | 1.80 ± 0.04 | 1.73 | 1.95 | 83 / 83 | b | -30% | < 0.0001 | | | |
| DAF-1 in set 2 | | | | | | | | | | | |
| | *daf-1(m40) IV (no ftEx205 transgene)* | 2.83 ± 0.09 | 2.85 | 3.20 | 61 / 61 | a | | | | | 3H |
| | *daf-1(m40) IV; ftEx205[Ptdc-1::daf-1-gfp + Podr-1::dsRed]* | 1.68 ± 0.04 | 1.71 | 1.86 | 97 / 97 | b | -41% | < 0.0001 | | | |
| DAF-1 in set 3 | | | | | | | | | | | |
| | *daf-1(m40) IV* (no *ftEx93* transgene) | 2.77 ± 0.06 | 2.88 | 3.31 | 136 / 136 | a | | | | | 3I |
| | *daf-1(m40) IV; ftEx93[Pglr-1::daf-1-gfp + Podr-1::dsRED]* | 1.84 ± 0.03 | 1.83 | 2.05 | 131 / 131 | b | -34% | < 0.0001 | | | |
| DAF-1 in set 4 | | | | | | | | | | | |
| | *daf-1(m40) IV* (no *ftEx74* transgene) | 2.47 ± 0.04 | 2.54 | 2.73 | 80 / 80 | a | | | | | S3D |
| | *daf-1(m40); ftEx74[Pglr-8p::daf-1-GFP + Podr-1::dsRED]* | 1.86 ± 0.03 | 1.84 | 2.04 | 103 / 103 | b | -25% | < 0.0001 | | | |
| DAF-1 in set 5 | | | | | | | | | | | |
| | *daf-1(m40) IV* (no *ftEx183* transgene) | 2.72 ± 0.06 | 2.75 | 3.19 | 99 / 99 | a | | | | | S3E |
| | *daf-1(m40) IV; ftEx183[Pglr-7::daf-1-gfp + Podr-1::dsRED]* | 2.68 ± 0.06 | 2.64 | 3.20 | 105 / 105 | b | -1% | > 0.05 | | | |
| DAF-3 in DAF-1 cells | | | | | | | | | | | |
| | *daf-1(m40) IV; daf-3(mgDf90) X* | 0.89 ± 0.04 | 0.93 | 1.02 | 35 / 35 | a | | | | | 3J |
| | *daf-1(m40) IV; daf-3(mgDf90) X; witEx4[Pdaf-1::daf-3 + Punc-122::RFP]* | 1.27 ± 0.04 | 1.31 | 1.42 | 47 / 47 | b | 43% | < 0.0001 | | | |
| DAF-3 in set 2 | | | | | | | | | | | |
| | *daf-1(m40) IV; daf-3(mgDf90) X* | 0.96 ± 0.03 | 0.89 | 1.03 | 42 / 42 | a | | | | | 3K |
| | *daf-1(m40) IV; daf-3(mgDf90) X; witEx2[Ptdc-1::daf-3 + Pmyo-2::RFP]* | 0.88 ± 0.04 | 0.85 | 1.09 | 54 / 54 | b | -8% | > 0.05 | | | |
| | *daf-1(m40) IV; daf-3(mgDf90) X* | 0.80 ± 0.02 | 0.78 | 0.89 | 38 / 40 | a | | | | | S3F |
| | *daf-1(m40) IV; daf-3(mgDf90) X; witEx3[Ptdc-1::daf-3 + Pmyo-2::RFP]* line 1 | 0.75 ± 0.04 | 0.67 | 0.86 | 25 / 26 | b | -4% | > 0.05 | | | |
| | *daf-1(m40) IV; daf-3(mgDf90) X; witEx2[Ptdc-1::daf-3 + Pmyo-2::RFP]* line 2 | 0.76 ± 0.03 | 0.72 | 0.85 | 38 / 40 | c | -4% | > 0.05 | | | |
| | *daf-1(m40) IV; daf-3(mgDf90) X; witEx1[Ptdc-1::daf-3 + Pmyo-2::RFP]* line 3 | 0.66 ± 0.04 | 0.68 | 0.76 | 26 / 26 | d | -17% | 0.02 | | | |

#### Table S4. Statistical analysis for Figures 4 and S4.

| Set | Genotype | Mean survival ± SEM (days) | Median survival (days) | 75th percentile (days) | N dead / initial N | Group | % Mean survival change vs. group a | *P* value (log-rank) vs. group a | *P* value (log-rank) vs. group b | *P* value (log-rank) vs. group c | *P* value (log-rank) vs. group e | Figure |
|---|---|---|---|---|---|---|---|---|---|---|---|---|
| Dauer | | | | | | | | | | | | |
| | wild type | 0.94 ± 0.02 | 0.91 | 1.09 | 118 / 118 | a | | | | | | 4B |
| | *daf-7(e1372) III* | 2.05 ± 0.07 | 1.91 | 2.41 | 81 / 81 | b | 118% | < 0.0001 | | | | |
| | *daf-12(rh61rh411) X* | 0.82 ± 0.02 | 0.78 | 0.91 | 59 / 59 | c | -13% | < 0.0001 | < 0.0001 | | | |
| | *daf-7(e1372) III; daf-12(rh61rh411) X* | 3.02 ± 0.09 | 3.04 | 3.64 | 82 / 82 | d | 222% | < 0.0001 | < 0.0001 | < 0.0001 | | |
| Fat storage | | | | | | | | | | | | |
| | wild type | 0.95 ± 0.01 | 0.92 | 1.08 | 235 / 235 | a | | | | | | 4C |
| | *daf-1(m40) IV* | 2.45 ± 0.05 | 2.34 | 2.87 | 193 / 200 | b | 158% | < 0.0001 | | | | |
| | *mgl-3(tm1766) IV; mgl-1(tm1811) X* | 0.98 ± 0.02 | 0.93 | 1.06 | 125 / 125 | c | 3% | > 0.05 | < 0.0001 | | | |
| | *daf-1(m40) mgl-3(tm1766) IV; mgl-1(tm1811) X* | 2.36 ± 0.04 | 2.36 | 2.79 | 226 / 226 | d | 148% | < 0.0001 | > 0.05 | < 0.0001 | | |
| | wild type | 0.96 ± 0.01 | 0.95 | 1.08 | 340 / 340 | a | | | | | | S4A,B |
| | *daf-1(m40) IV* | 2.44 ± 0.04 | 2.40 | 2.87 | 291 / 298 | b | 155% | < 0.0001 | | | | |
| | *mgl-1(tm1811) X* | 0.85 ± 0.02 | 0.84 | 0.93 | 101 / 101 | c | -11% | < 0.0001 | < 0.0001 | | | |
| | *mgl-1(tm1811) X; daf-1(m40) IV* | 2.33 ± 0.04 | 2.37 | 2.60 | 94 / 94 | d | 143% | < 0.0001 | 0.001 | < 0.0001 | | |
| | *mgl-3(tm1766) IV* | 1.17 ± 0.02 | 1.17 | 1.33 | 98 / 103 | e | 22% | < 0.0001 | < 0.0001 | | | |
| | *daf-1(m40) mgl-3(tm1766) IV* | 2.39 ± 0.06 | 2.35 | 2.86 | 100 / 100 | f | 149% | < 0.0001 | > 0.05 | | < 0.0001 | |
| Germline | | | | | | | | | | | | |
| | *mes-1(ok2467) X* (fertile) | 0.97 ± 0.03 | 0.92 | 1.10 | 97 / 97 | a | | | | | | 4D |
| | *daf-1(m40) IV; mes-1(ok2467) X* (fertile) | 2.39 ± 0.05 | 2.44 | 2.67 | 85 / 85 | b | 147% | < 0.0001 | | | | |
| | *mes-1(ok2467) X* (sterile) | 1.52 ± 0.04 | 1.55 | 1.80 | 105 / 105 | c | 57% | < 0.0001 | < 0.0001 | | | |
| | *daf-1(m40) IV; mes-1(ok2467) X* (sterile) | 2.89 ± 0.12 | 2.94 | 3.57 | 52 / 52 | d | 198% | < 0.0001 | < 0.0001 | < 0.0001 | | |
| | *mes-1(ok2467) X* (fertile) | 0.93 ± 0.02 | 0.91 | 1.05 | 123 / 123 | a | | | | | | 4E |
| | *daf-3(mgDf90) mes-1(ok2467) X* (fertile) | 0.96 ± 0.02 | 0.96 | 1.09 | 126 / 126 | b | 3% | > 0.05 | | | | |
| | *mes-1(ok2467) X* (sterile) | 1.44 ± 0.03 | 1.49 | 1.67 | 126 / 126 | c | 54% | < 0.0001 | < 0.0001 | | | |
| | *daf-3(mgDf90) mes-1(ok2467) X* (sterile) | 1.36 ± 0.03 | 1.36 | 1.53 | 123 / 123 | d | 45% | < 0.0001 | < 0.0001 | > 0.05 | | |

### Table S5. Statistical analysis for Figures 5 and S5.

| Set | Genotype | Mean survival ± SEM (days) | Median survival (days) | 75th percentile (days) | N dead / initial N | Group | % Mean survival change vs. group a | *P* value (log-rank) vs. group a | *P* value (log-rank) vs. group b | *P* value (log-rank) vs. group c | Figure |
|---|---|---|---|---|---|---|---|---|---|---|---|
| Pathway analysis | | | | | | | | | | | |
| | wild type | 0.89 ± 0.01 | 0.87 | 1.03 | 275 / 275 | a | | | | | |
| | *ins-4(tm3620)* | 1.16 ± 0.02 | 1.18 | 1.32 | 120 / 120 | b | 30% | < 0.0001 | | | |
| | *ins-5(tm2560) II* | 1.09 ± 0.03 | 1.11 | 1.28 | 86 / 86 | c | 22% | < 0.0001 | | | 5A |
| | *ins-6(tm2416) II* | 1.47 ± 0.03 | 1.49 | 1.69 | 90 / 90 | d | 65% | < 0.0001 | | | |
| | *daf-28(tm2308) V* | 1.18 ± 0.02 | 1.15 | 1.30 | 137 / 137 | e | 32% | < 0.0001 | | | |
| | wild type | 0.77 ± 0.01 | 0.74 | 0.83 | 128 / 128 | a | | | | | S5A |
| | *ins-1(nr2091) IV* | 0.95 ± 0.02 | 0.94 | 1.04 | 113 / 113 | b | 24% | < 0.0001 | | | |
| | wild type | 0.98 ± 0.02 | 0.96 | 1.10 | 147 / 147 | a | | | | | S5B |
| | *ins-3(tm3608) II* | 1.08 ± 0.02 | 1.10 | 1.23 | 141 / 141 | b | 11% | < 0.0001 | | | |
| | wild type | 1.03 ± 0.01 | 1.03 | 1.12 | 129 / 140 | a | | | | | |
| | *daf-1(m40) IV* | 2.79 ± 0.08 | 2.76 | 3.63 | 160 / 183 | b | 171% | < 0.0001 | | | 5B |
| | *daf-2(e1370) III* | 3.02 ± 0.06 | 2.87 | 3.57 | 166 / 185 | c | 193% | < 0.0001 | > 0.05 | | |
| | *daf-2(e1370) III; daf-1(m40) IV* | 5.88 ± 0.21 | 5.63 | 7.12 | 77 / 159 | d | 471% | < 0.0001 | < 0.0001 | < 0.0001 | |
| | wild type | 0.84 ± 0.01 | 0.82 | 0.92 | 145 / 158 | a | | | | | |
| | *daf-2(e1370) III* | 3.50 ± 0.06 | 3.46 | 4.07 | 136 / 154 | b | 318% | < 0.0001 | | | 5C |
| | *daf-16(mu86) I* | 0.78 ± 0.01 | 0.76 | 0.86 | 141 / 153 | c | -7% | 0.004 | < 0.0001 | | |
| | *daf-16(mu86) I; daf-2(e1370) III* | 0.72 ± 0.01 | 0.71 | 0.79 | 140 / 151 | d | -14% | < 0.0001 | < 0.0001 | < 0.0001 | |
| | *ayIs7[Phlh-8::GFP + dpy-20(+)]* | 0.83 ± 0.01 | 0.84 | 0.92 | 145 / 145 | a | | | | | |
| | *ins-4 ins-5 ins-6(hpDf761) II; ayIs7 IV; daf-28(tm2308) V* | 1.26 ± 0.02 | 1.25 | 1.45 | 131 / 131 | b | 52% | < 0.0001 | | | 5D |
| | *daf-16(mgDf50) I* | 0.77 ± 0.01 | 0.73 | 0.83 | 133 / 133 | c | -8% | 0.012 | < 0.0001 | | |
| | *daf-16(mgDf50) I; ins-4 ins-5 ins-6(hpDf761) II; ayIs7 IV; daf-28(tm2308) V* | 0.69 ± 0.01 | 0.69 | 0.78 | 140 / 140 | d | -17% | < 0.0001 | < 0.0001 | < 0.0001 | |
| | wild type | 0.89 ± 0.02 | 0.86 | 0.98 | 159 / 161 | a | | | | | |
| | *daf-7(e1372) III* | 2.46 ± 0.06 | 2.44 | 2.91 | 114 / 118 | b | 175% | < 0.0001 | | | 5E |
| | *daf-16(mu86) I* | 0.68 ± 0.01 | 0.67 | 0.73 | 163 / 165 | c | -24% | < 0.0001 | < 0.0001 | | |
| | *daf-16(mu86) I; daf-7(e1372) III* | 1.16 ± 0.02 | 1.15 | 1.30 | 160 / 168 | d | 29% | < 0.0001 | < 0.0001 | < 0.0001 | |
| | wild type | 0.99 ± 0.01 | 0.98 | 1.09 | 143 / 143 | a | | | | | |
| | *daf-1(m40) IV* | 2.45 ± 0.05 | 2.57 | 2.79 | 127 / 127 | b | 147% | < 0.0001 | | | 5F |
| | *daf-16(mu86) I* | 0.84 ± 0.01 | 0.83 | 0.93 | 105 / 105 | c | -15% | < 0.0001 | < 0.0001 | | |
| | *daf-16(mu86) I; daf-1(m40) IV* | 1.39 ± 0.02 | 1.43 | 1.51 | 123 / 123 | d | 40% | < 0.0001 | < 0.0001 | < 0.0001 | |
| | *daf-1(m40) IV* | 2.79 ± 0.08 | 2.78 | 3.50 | 98 / 98 | a | | | | | S5C |
| | *daf-1(m40) hlh-30(tm1978) IV* | 2.68 ± 0.06 | 2.65 | 3.04 | 121 / 121 | b | -4% | 0.04 | | | |
| | wild type + empty vector | 0.90 ± 0.02 | 0.85 | 1.01 | 123 / 123 | a | | | | | |
| | *daf-1(m40) IV* + empty vector | 2.15 ± 0.06 | 2.17 | 2.46 | 109 / 109 | b | 140% | < 0.0001 | | | 5G |
| | *skn-1(RNAi)* | 0.91 ± 0.01 | 0.92 | 1.01 | 130 / 130 | c | 2% | > 0.05 | < 0.0001 | | |
| | *daf-1(m40) IV + skn-1(RNAi)* | 1.52 ± 0.04 | 1.53 | 1.80 | 107 / 107 | d | 69% | < 0.0001 | < 0.0001 | < 0.0001 | |
| | *daf-16(mu86) I* + empty vector | 0.77 ± 0.02 | 0.77 | 0.89 | 119 / 119 | a | | | | | |
| | *daf-16(mu86) I; daf-1(m40) IV* + empty vector | 1.26 ± 0.03 | 1.26 | 1.46 | 92 / 92 | b | 64% | < 0.0001 | | | 5H |
| | *daf-16(mu86) I + skn-1(RNAi)* | 0.57 ± 0.01 | 0.56 | 0.64 | 134 / 134 | c | -26% | < 0.0001 | < 0.0001 | | |
| | *daf-16(mu86) I; daf-1(m40) IV + skn-1(RNAi)* | 0.79 ± 0.02 | 0.82 | 1.00 | 114 / 114 | d | 3% | > 0.05 | < 0.0001 | < 0.0001 | |
| Full DAF-16 expression | | | | | | | | | | | |
| | *daf-16(mu86) I; daf-1(m40) IV* | 1.44 ± 0.02 | 1.48 | 1.59 | 130 / 130 | a | | | | | |
| | *daf-16(mu86) I; daf-1(m40) IV; qyIs288[Pdaf-16::GFP::daf-16 + unc-119(+)]* | 2.62 ± 0.07 | 2.66 | 3.20 | 120 / 120 | b | 82% | < 0.0001 | | | 5I |
| | *daf-1(m40) IV* | 2.38 ± 0.04 | 2.43 | 2.63 | 88 / 88 | c | 65% | < 0.0001 | < 0.0001 | | |
| DAF-16 in intestine | | | | | | | | | | | |
| | *daf-16(mu86) I; daf-1(m40) IV* | 1.65 ± 0.02 | 1.68 | 1.78 | 71 / 74 | a | | | | | |
| | *daf-16(mu86) I; daf-1(m40) IV; qyIs292[Pges-1::GFP::daf-16]* | 2.22 ± 0.03 | 2.17 | 2.47 | 93 / 97 | b | 34% | < 0.0001 | | | 5J |
| | *daf-1(m40) IV* | 2.72 ± 0.06 | 2.74 | 3.14 | 61 / 72 | c | 64% | < 0.0001 | < 0.0001 | | |
| DAF-16 in neurons | | | | | | | | | | | |
| | *daf-16(mu86) I; daf-1(m40) IV* | 1.63 ± 0.02 | 1.63 | 1.80 | 130 / 130 | a | | | | | |
| | *daf-16(mu86) I; daf-1(m40) IV; qyIs294[Punc-119::GFP::daf-16]* | 1.76 ± 0.03 | 1.76 | 1.99 | 134 / 134 | b | 8% | 0.0004 | | | 5K |
| | *daf-1(m40) IV* | 2.35 ± 0.06 | 2.42 | 2.82 | 124 / 124 | c | 44% | < 0.0001 | < 0.0001 | | |
| DAF-16 in body muscle | | | | | | | | | | | |
| | *daf-16(mu86) I; daf-1(m40) IV* | 1.47 ± 0.02 | 1.48 | 1.61 | 113 / 113 | a | | | | | |
| | *daf-16(mu86) I; unc-119(ed4) III; daf-1(m40) IV; qyEx264 [Pmyo-3::GFP::daf-16 + unc-119(+)]* | 1.47 ± 0.03 | 1.49 | 1.63 | 114 / 114 | b | 0% | > 0.05 | | | 5L |
| | *daf-1(m40) IV* | 2.38 ± 0.06 | 2.45 | 2.91 | 114 / 114 | c | 62% | < 0.0001 | < 0.0001 | | |
| DAF-16 in hypodermis | | | | | | | | | | | |
| | *daf-16(mu86) I; daf-1(m40) IV* | 1.52 ± 0.03 | 1.55 | 1.66 | 49 / 49 | a | | | | | |
| | *daf-16(mu86) I; daf-1(m40) IV; qyIs290[Pcol-12::GFP::daf-16 + unc-119(+)]* | 1.13 ± 0.09 | 1.25 | 1.50 | 29 / 29 | b | -26% | 0.008 | | | 5M |
| | *daf-1(m40) IV* | 2.78 ± 0.05 | 2.78 | 3.12 | 82 / 82 | c | 83% | < 0.0001 | < 0.0001 | | |

#### Table S6. Statistical analysis for Figures 6 and S6.

| Set | Genotype or condition | Mean survival ± SEM (days) | Median survival (days) | 75th percentile (days) | N dead / initial N | Group | % Mean survival change vs. group a | *P* value (log-rank) vs. group a | *P* value (log-rank) vs. group b | *P* value (log-rank) vs. group c | Figure |
|---|---|---|---|---|---|---|---|---|---|---|---|
| Pathway analysis | | | | | | | | | | | |
| | wild type | 0.89 ± 0.02 | 0.87 | 1.00 | 139 / 139 | a | | | | | 6A |
| | *daf-1(m40) IV* | 2.26 ± 0.05 | 2.28 | 2.67 | 125 / 125 | b | 153% | < 0.0001 | | | |
| | *tbh-1(ok1196) X* | 0.91 ± 0.01 | 0.87 | 1.02 | 139 / 139 | c | 2% | > 0.05 | < 0.0001 | | |
| | *daf-1(m40) IV; tbh-1(ok1196) X* | 2.83 ± 0.04 | 2.83 | 3.10 | 142 / 142 | d | 217% | < 0.0001 | < 0.0001 | < 0.0001 | |
| | *daf-1(m40) IV* | 1.79 ± 0.05 | 1.73 | 2.11 | 94 / 94 | a | | | | | S6A |
| | *tdc-1(ok914) II; daf-1(m40) IV* | 2.72 ± 0.05 | 2.62 | 3.05 | 128 / 128 | b | 52% | < 0.0001 | | | |
| 1-day food-level conditioning | | | | | | | | | | | |
| | 250 x $10^6$ *E. coli* OP50 lawn | 1.08 ± 0.02 | 1.09 | 1.24 | 107 / 121 | a | | | | | 6B |
| | 50 x $10^6$ *E. coli* OP50 lawn | 1.06 ± 0.02 | 1.09 | 1.18 | 113 / 115 | b | -2% | > 0.05 | | | |
| | 10 x $10^6$ *E. coli* OP50 lawn | 0.93 ± 0.03 | 0.96 | 1.09 | 71 / 77 | c | -14% | 0.009 | 0.0009 | | |
| | 0 *E. coli* OP50 lawn | 0.29 ± 0.02 | 0.30 | 0.37 | 56 / 56 | d | -73% | < 0.0001 | < 0.0001 | < 0.0001 | |
| 2-day food-level conditioning | | | | | | | | | | | |
| | 200 x $10^6$ *E. coli* OP50 lawn | 0.54 ± 0.01 | 0.52 | 0.61 | 141 / 148 | a | | | | | S6B |
| | 100 x $10^6$ *E. coli* OP50 lawn | 0.56 ± 0.02 | 0.51 | 0.66 | 139 / 148 | b | 3% | > 0.05 | | | |
| | 50 x $10^6$ *E. coli* OP50 lawn | 0.48 ± 0.02 | 0.43 | 0.57 | 144 / 149 | c | -12% | 0.03 | 0.0009 | | |
| | 25 x $10^6$ *E. coli* OP50 lawn | 0.21 ± 0.01 | 0.19 | 0.24 | 159 / 162 | d | -62% | < 0.0001 | < 0.0001 | < 0.0001 | |
| | with food | 0.73 ± 0.01 | 0.71 | 0.81 | 123 / 141 | a | | | | | 6C |
| | no food | 0.12 ± 0.00 | 0.12 | 0.13 | 97 / 97 | b | -84% | < 0.0001 | | | |
| Food ingestion | | | | | | | | | | | |
| | wild type | 0.81 ± 0.01 | 0.79 | 0.93 | 152 / 152 | a | | | | | 6D |
| | *eat-2(ad1116) II* | 0.62 ± 0.01 | 0.61 | 0.68 | 152 / 153 | b | -23% | < 0.0001 | | | |
| | *daf-3(mgDf90) X* | 0.84 ± 0.01 | 0.83 | 0.93 | 118 / 118 | c | 4% | > 0.05 | < 0.0001 | | |
| | *eat-2(ad1116) II; daf-3(mgDf90) X* | 0.45 ± 0.01 | 0.45 | 0.49 | 128 / 128 | d | -44% | < 0.0001 | < 0.0001 | < 0.0001 | |

### Table S7. Statistical analysis for Figures 7 and S7.

| Set | Worm genotype | *E. coli* genotype | Mean survival ± SEM (days) | Median survival (days) | 75th percentile (days) | N dead / initial N | Group | % Mean survival change vs. group a | *P* value (log-rank) vs. group a | *P* value (log-rank) vs. group b | *P* value (log-rank) vs. group c | Figure |
|---|---|---|---|---|---|---|---|---|---|---|---|---|
| Bacterial genotype, 1 mM $H_2O_2$ | | | | | | | | | | | | |
| | wild type | *katG katE ahpCF* | 1.42 ± 0.06 | 1.26 | 1.49 | 53 / 53 | a | | | | | 7B |
| | *daf-7(ok3125) III* | *katG katE ahpCF* | 3.30 ± 0.21 | 3.25 | 3.46 | 25 / 25 | b | 132% | < 0.0001 | | | |
| | wild type | wild type | > 6.36 | . | . | 2 / 78 | c | > 347% | < 0.0001 | < 0.0001 | | |
| | *daf-7(ok3125) III* | wild type | > 5.74 | . | . | 3 / 80 | d | > 304% | < 0.0001 | < 0.0001 | > 0.05 | |
| Pathway analysis | | | | | | | | | | | | |
| | wild type | *katG katE ahpCF* | 1.97 ± 0.04 | 1.96 | 2.31 | 137 / 149 | a | | | | | S7A |
| | ASI: ablation | *katG katE ahpCF* | 2.57 ± 0.05 | 2.55 | 2.98 | 122 / 138 | b | 31% | < 0.0001 | | | |
| | wild type | *katG katE ahpCF* | 1.91 ± 0.06 | 1.74 | 2.43 | 126 / 126 | a | | | | | 7C |
| | *wuls151[ctl-1(+) + ctl-2(+) + ctl-3(+) + Pmro-3::GFP]* | *katG katE ahpCF* | 5.12 ± 0.19 | 5.18 | 6.45 | 96 / 96 | b | 168% | < 0.0001 | | | |
| | wild type | *katG katE ahpCF* | 1.14 ± 0.03 | 1.06 | 1.37 | 121 / 121 | a | | | | | 7E |
| | *daf-1(m40) IV* | *katG katE ahpCF* | 2.70 ± 0.10 | 2.64 | 3.35 | 85 / 85 | b | 138% | < 0.0001 | | | |
| | *ctl-1(ok1242) II* | *katG katE ahpCF* | 0.91 ± 0.04 | 0.87 | 1.11 | 93 / 93 | c | -20% | 0.0002 | < 0.0001 | | |
| | *ctl-1(ok1242) II; daf-1(m40) IV* | *katG katE ahpCF* | 1.51 ± 0.08 | 1.44 | 1.94 | 97 / 97 | d | 33% | < 0.0001 | < 0.0001 | < 0.0001 | |
| | wild type | *katG katE ahpCF* | 0.84 ± 0.03 | 0.81 | 1.01 | 110 / 110 | a | | | | | S7B |
| | *daf-1(m40) IV* | *katG katE ahpCF* | 4.26 ± 0.19 | 3.78 | 5.19 | 109 / 109 | b | 408% | < 0.0001 | | | |
| | *ctl-2(ok1137) II* | *katG katE ahpCF* | 1.24 ± 0.04 | 1.16 | 1.43 | 112 / 112 | c | 48% | < 0.0001 | < 0.0001 | | |
| | *ctl-2(ok1137) II; daf-1(m40) IV* | *katG katE ahpCF* | 4.24 ± 0.19 | 3.78 | 5.26 | 115 / 115 | d | 405% | < 0.0001 | > 0.05 | < 0.0001 | |

#### Table S8: Bacterial strains.

| Strain | Genotype | Source | Reference |
|---|---|---|---|
| OP50 | *E. coli* B, uracil auxotroph | CGC | Brenner, 1974 |
| HT115 | *E. coli* $F^-$ *mcrA, mcrB, IN(rrnD-rrnE)1, rnc14::Tn10(DE3 lysogen: lavUV5 promoter -T7 polymerase).* Tetracycline resistant | CGC | Kamath et al., 2001 |
| MG1655 | *E. coli* K12 $F^-$ wild type | James Imlay | Seaver and Imlay, 2001 |
| JI377 | *E. coli* MG1655 *ahpCF katG katE* | James Imlay | Seaver and Imlay, 2001 |

### Table S9: *C. elegans* strains.

| Strain | Genotype | Comment | Figure | Source | Reference |
|---|---|---|---|---|---|
| PR813 | osm-5(p813) X | | 1 | CGC | |
| PR671 | tax-2(p671) I | | S1 | CGC | |
| PY7512 | tax-4(p678) III; Ex[Pttx-1::tax-4 + Punc-122::dsred] | | 1 | Piali Sengupta | |
| FG540 | udEx428[Psrh-124::ced- 3(p15), Psrh-142::ced- 3(p17), Psrh-142::gfp, Pelt-2::gfp] | ADF ablation | 1 | Denise Ferkey | Krzyzanowski et al., 2016 |
| HCX179 | bosEx179[Psre-1::csp-1b + Punc-122::GFP] | ADL ablation | S1 | Howard Chang | Horspool and Chang, 2017 |
| SAY141 | pgIs2[Pgcy-8::TU#813 + Pgcy-8::TU#814 + Punc-122::GFP + Pgcy-8::mCherry + Pgcy-8::GFP + Pttx-3::GFP] | AFD ablation | 1 | CGC, This work | Glauser et al., 2011 |
| JPS271 | vxEx265 [gcy-8p::ICE + myo-2p:mCherry]. | AFD ablation | S1 | CGC | Vidal-Gadea et al., 2015 |
| OH13098 | che-1(ot75) I | ASE fate determinant | 1 | CGC | Chang et al., 2003 |
| RJP3130 | otEx3130Pops-1::p12 caspase + Pgcy-21 p17 caspase + Pmyo-3::RFP]; Is[Pets-5::mCherry + Pelt-2::GFP] | ASG ablation | 1 | Roger Pocock | Juozaityte et al., 2017 |
| SAY113 | Is[Psra-6::mCasp1 + Pmyo-3::RFP] | ASH ablation | 1 | Takaaki Hirotsu | Yoshida et al., 2012 |
| PY7505 | oyIs84 [Pgpa-4::TU#813 + Pgcy-27::TU#814 + Pgcy-27::GFP + Punc-122::DsRed] | ASI ablation | 1, S7 | CGC | Beverly et al., 2011 |
| ZD763 | mgIs40[Pdaf-28::nls-GFP]; jxEx102[Ptrx-1::ICE + Pofm-1::gfp] | ASJ ablation | 1 | Dennis Kim | Cornils et al., 2011 |
| SAY119 | qrIs2[Psra-9::mCasp1] 2x outcrossed | ASK ablation | 1 | CGC, This work | Srinivasan et al., 2012 |
| SAY114 | Ex[odr-10::mCasp1 + myo-3::GFP] | AWA ablation | 1 | Takaaki Hirotsu | Yoshida et al., 2012 |
| SAY115 | Is[str-1::mCasp1 + myo-3::GFP ] | AWB ablation | 1 | Takaaki Hirotsu | Yoshida et al., 2012 |
| PY7502 | oyIs85[Pceh-36::TU#813 + Pceh-36::TU#814 + Psrtx-1::GFP + Punc-122::dsRed] | AWC ablation | 1 | CGC | Beverly et al., 2011 |
| LJ200 | unc-119(ed3); ysIs1[punc-119cR + Pklp-6::ced-3 + Pklp-6:egl-1] | IL2 ablation | 1 | Junho Lee | Lee et al., 2011 |
| ZD653 | qdEx22[Pser-2::csp-1b; Pmyo-2::rfp] | OLL ablation | 1 | Dennis Kim | Chang et al., 2011 |
| VM6365 | lin-15(n765ts) X; akEx387[lin-15(+) + Pdat-1::gfp + Pdat-1::ICE] | ADE/PDE/CEP ablation | 1 | Andres Maricq | Wragg et al., 2007 |
| JPS278 | vxEx277[Pmec-3::ICE + Pmyo-2::mCherry] | ALM/PLM/AVM/PVM/FLP/PVD ablation | 1 | CGC | Russell et al., 2014 |
| CX7102 | lin-15B&lin-15A(n765) qaIs2241[Pgcy-36::egl-1 + Pgcy-35::GFP + lin-15(+)] X | URX/AQR/PQR ablation | 1 | CGC | Chang et al., 2006 |
| RB2302 | daf-7(ok3125) III | | 2,7, S2-S3 | CGC | |
| ZD729 | daf-7(ok3125) III; qdEx37[Pdaf-7::daf-7 + Pges-1::GFP] | | 2 | Dennis Kim | Fletcher and Kim, 2017 |
| ZD732 | daf-7(ok3125) III; qdEx40[Ptrx-1::daf-7 + Pges-1::GFP] | | 2 | Dennis Kim | Fletcher and Kim, 2017 |
| ZD736 | daf-7(ok3125) III; qdEx44[Pstr-3::daf-7 + Pges-1::GFP] | | 2 | Dennis Kim | Fletcher and Kim, 2017 |
| GR1311 | daf-3(mgDf90) X | | 3 | CGC | |
| SAY84 | daf-3(mgDf90) X; oyIs84[Pgpa-4::TU#813 + Pgcy-27::TU#814 + Pgcy-27::GFP + Punc-122::DsRed] | | 3 | This work | |
| SAY85 | daf-1(m40) IV 6x outcrossed | | 2-7, S2-S3, S7 | This work | |
| SAY72 | daf-1(m40) IV; daf-3(mgDf90) X | | 3, S3 | This work | |
| CB1372 | daf-7(e1372) III | | 5, S2 | CGC | |
| CB1376 | daf-3(e1376) X | | S3 | CGC | |
| ZD907 | daf-7(ok3125) III; daf-3(e1376) X | | S3 | Dennis Kim | Fletcher and Kim, 2017 |
| CB1386 | daf-5(e1386) II | | S3 | CGC | |
| GR1278 | daf-5(e1386) II; daf-7(e1372) III | | S3 | Jane Hubbard | Dalfo et al., 2012 |
| GC1149 | daf-5(e1386) II; daf-1(m40) IV | | S3 | Jane Hubbard | Dalfo et al., 2012 |
| KQ275 | daf-1(m40) IV; ftEx93[pglr-1::daf-1-gfp odr-1::dsRED] | | 3 | Young-Jai You | Greer et al., 2008 |
| KQ315 | daf-1(m40) IV; ftEx166[pflp-1::daf-1::gfp + podr-1::dsRED] | | 3 | Young-Jai You | Greer et al., 2008 |
| KQ280 | daf-1(m40) IV; ftEx98[Pdaf-1::daf-1-gfp + Podr-1::dsRED] | | 3 | Dennis Kim | Greer et al., 2008 |
| KQ251 | daf-1(m40) IV; ftEx69[Pegl-3:: daf-1-gfp + Podr-1::dsRED] | | 3 | Dennis Kim | Greer et al., 2008 |
| KQ380 | daf-1(m40) IV; ftEx205[Ptdc-1::daf-1-gfp + Podr-1::dsRED] | | 3 | Dennis Kim | Greer et al., 2008 |
| KQ265 | daf-1(m40) IV; ftEx83[Posm-6::daf-1::GFP + Podr-1::dsRED] | | 3 | Dennis Kim | Greer et al., 2008 |
| KQ332 | daf-1(m40) IV; ftEx183[pglr-7::daf-1-gfp odr-1::dsRED] | | S3 | Young-Jai You | Greer et al., 2008 |
| KQ256 | daf-1(m40) IV; ftEx83[Pglr-8::daf-1::GFP + Podr-1::dsRED] | | S3 | Dennis Kim | Greer et al., 2008 |
| SAY137 | daf-1(m40) IV; daf-3(mgDf90) X; witEx4[Pdaf-1::daf-3(+)-GFP + Punc-122::dsRED] | | 3 | This work | |
| SAY118 | daf-1(m40) IV; daf-3(mgDf90) X; witEx1[Ptdc-1::daf-3(+)::GFP+ Pmyo-2::RFP] | | S3 | This work | |
| SAY133 | daf-1(m40) IV; daf-3(mgDf90) X; witEx2[Ptdc-1::daf-3(+)::GFP + Pmyo-2::RFP] | | 3 | This work | |
| SAY134 | daf-1(m40) IV; daf-3(mgDf90) X; witEx3[Ptdc-1::daf-3(+)::GFP + Pmyo-2::RFP] | | S3 | This work | |
| QZ117 | daf-12(rh61rh411) X | | 4 | Joy Alcedo | |
| QC1239 | daf-7(e1372) III; daf-12(rh61rh411) X | | 4 | Jane Hubbard | Dalfo et al., 2012 |
| ZD1424 | daf-1(m40) mgl-3(tm1766) IV; mgl-1(tm1811) X | | 4 | Dennis Kim | Fletcher and Kim, 2017 |
| SAY92 | mgl-3(tm1766) IV; mgl-1(tm1811) X | | 4 | This work | |
| SAY124 | mgl-1(tm1811)X; daf-1(m40)IV | | S4 | This work | |
| SAY130 | mgl-1(tm1811) X | | S4 | This work | |
| SAY128 | mgl-3(tm1766) daf-1(m40) IV | | S4 | This work | |
| SAY129 | mgl-3(tm1766) IV | | S4 | This work | |
| SAY80 | daf-1(m40) IV; mes-1(ok2467) X | | 4 | This work | |
| SAY20 | mes-1(ok2467) X 6x outcrossed | | 4 | This work | |
| SAY110 | daf-3(mgDf90) mes-1(ok2467) X | | 4 | This work | |
| QZ91 | daf-2(e1370) III 6x outcrossed | | 5 | Joy Alcedo | Fernandes de Abreu et al., 2014 |
| SAY77 | daf-2(e1370) III; daf-1(m40) IV | | 5 | This work | |
| QZ80 | ins-1(nr2091) IV | | S5 | Yun Zhang | Fernandes de Abreu et al., 2014 |
| QL28 | ins-3(tm3608) II | | S5 | Yun Zhang | Fernandes de Abreu et al., 2014 |
| QL27 | ins-4(tm3620) II | | 5 | Yun Zhang | Fernandes de Abreu et al., 2014 |
| QL24 | ins-5(tm2560) II | | 5 | Yun Zhang | Fernandes de Abreu et al., 2014 |
| QZ81 | ins-6(tm2416) II | | 5 | Yun Zhang | Fernandes de Abreu et al., 2014 |
| QZ83 | daf-28(tm2308) V | | 5 | Yun Zhang | Fernandes de Abreu et al., 2014 |
| QZ60 | daf-16(mu86) I | | 5 | Joy Alcedo | Fernandes de Abreu et al., 2014 |
| QZ218 | daf-16(mu86) I; daf-2(e1370) III | | 5 | Joy Alcedo | Fernandes de Abreu et al., 2014 |
| GR1307 | daf-16(mgDf50) I | | 5 | CGC | |
| LRB209 | ins-4 ins-5 ins-6(hpDf761) II; ayIs7 IV; daf-28(tm2308) V | | 5 | Ryan Baugh | Kaplan et al., 2015 |
| LRB228 | daf-16(mgDf50) I; ins-4 ins-5 ins-6(hpDf761) II; ayIs7 IV; daf-28(tm2308) V | | 5 | Ryan Baugh | Kaplan et al., 2015 |
| GC1248 | daf-16(mu86) I; daf-7(e1372) III | | 5 | Jane Hubbard | Dalfo et al., 2012 |
| SAY74 | daf-16(mu86) I; daf-1(m40) IV | | 5 | This work | |
| SAY186 | daf-1(m40) hlh-30(tm1978) IV | | S5 | This work | |
| SAY102 | daf-16(mu86) I; daf-1(m40) IV; qyIs288[Pdaf-16::GFP::daf-16 + Punc-119(+)] | | 5 | This work | |
| SAY103 | daf-16(mu86) I; daf-1(m40) IV; unc-119(ed4) III; qyEx264[Pmyo-3::GFP::daf-16a + unc-119(+)] | | 5 | This work | |
| SAY104 | daf-16(mu86) I; daf-1(m40) IV; qyIs290[Pcol-12::GFP::daf-16 + unc-119(+)] | | 5 | This work | |
| SAY105 | daf-16(mu86) I; daf-1(m40) IV; qyIs292[Pges-1::GFP::daf-16] | | 5 | This work | |
| SAY100 | daf-16(mu86) I; daf-1(m40) IV; qyIs294[Punc-119::GFP::daf-16] | | 5 | This work | |
| SAY65 | tbh-1(ok1196) X | | 6 | This work | |
| KQ364 | daf-1(m40) IV; tbh-1(ok1196) X | | 6 | Dennis Kim | Fletcher and Kim, 2017 |
| SAY64 | daf-1(m40) IV; tdc-1(ok914) II | | S6 | Dennis Kim | Fletcher and Kim, 2017 |
| QZ414 | eat-2(ad1116) II | | 6 | Joy Alcedo | |
| SAY89 | eat-2(ad1116) II; daf-3(mgDf90) X | | 6 | This work | |
| GA800 | wuIs151[ctl-1(+) + ctl-2(+) + ctl-3(+) + Pmro-3::GFP] | | 7 | CGC | |
| UN1781 | ctl-1(ok1242) II | | 7 | Erin Cram | |
| SAY200 | ctl-1(ok1242) II; daf-1(m40) IV | | 7 | This work | |
| UN18129 | ctl-2(ok1137) II | | S7 | Erin Cram | |
| SAY191 | ctl-2(ok1137) II; daf-1(m40) IV | | S7 | This work | |
| KHA166 | chuIs166[unc-119(+), Pctl-1::Bxy-ctl-1::gfp] | | 7 | Koichi Hasegawa | Hamaguchi et al., 2019 |
| SAY231 | daf-1(m40) IV; chuIs166[unc-119(+), Pctl-1::Bxy-ctl-1::gfp] | | 7 | This work | |

**Table S10: PCR genotyping primers and enzymes, and phenotypes used for strain construction.**

| Allele | phenotype | Primers | Restriction Enzyme |
|---|---|---|---|
| *daf-3(mgDf90)* | - | AAACGCGTCATGTGGACCAC<br>ACCCTCATGCCTACTGTCAG | - |
| *oyIs84* | fluorescent marker | - | - |
| *daf-1(m40)* | dauer constitutive | GAGACGATCATCCACTTGGTAAG<br>AAATCTTCCGGATCCAACTCTAC | Tsp45I |
| *daf-7(ok3125)* | dauer constitutive | ATCTTCACTCCCGGTGTTTATCT<br>CTCAGGATTGGAGACTTTGTGAG | - |
| *mes-1(ok2467)* | sterile | - | - |
| *tdc-1(ok914)* | - | AACGGTGCATTTTTCAGGAC<br>GGACGTTGAGAATGCGAAAT<br>AAATGGTTTACGGGCTTGG<br>ATGGTTGGCCATGTTGAGAT | - |
| *eat-2(ad1116)* | decreased pumping | GGAGCCACTTAGGACACC<br>CCACACTATCTTTCTACCAC | Hpy188I |
| *tbh-1(ok1196)* | - | AAGCAGGATCAGGAGCACAT<br>ATGAGAAGTGCCGTTGCTCT<br>CATGTCATTGATGGCTGGAC<br>GAACGCCAGTTGGTTGATTT | - |
| *daf-2(e1370)* | dauer constitutive | GACGATCCCGAGGTGAGTAT<br>CAGCGATGGTTGTGATGGAA | - |
| *daf-16(mu86)* | - | AGAACACCATGGGGGCACTGGAT<br>GGCGGGAATGAAGCAAGAGCCAA<br>TGACGCTCACCTTGAAAAGGTCAAT<br>GGAACCGATTCGCCAACCCATGA | - |
| *ins-3(tm3608)* | - | GATCAGCATTGTCACCTGAC<br>TGGCAACTGATGTCCGGTAT | - |
| *ins-4(tm3620)* | - | CCGCCCAATCCCTTTAACGT<br>GATGGCTTGTTGGACGACTG<br>TGGCGCTTGACGCATCAGTC<br>GAGTGCACTGTGTTGTGCAG | - |
| *ins-5(tm2560)* | - | GGCTCCTTGCGCCATGATGT<br>ATCCTTGAATGCCCGTGAGT<br>GCTCCTCCCATGTTGGATTG<br>GGTTTCAGGAGGGTGACGGT | - |
| *ins-6(tm2416)* | - | TTTACCCACCCCTTCGTGAT<br>TTGTACAAGCCACTGGGATG<br>TCGTGATGCTCCGCCTATTG<br>ACAGAGACTGATATCGGAGT | - |
| *daf-28(tm2308)* | - | TCCGCCCACTTTGAGCTATA<br>GCACCCGATCTGACGACACT<br>GGGTTATCACTAGGAAGTTG<br>ACCGAGAGGTAGGGGTAATT | - |
| *hlh-30(tm1978)* | - | TGTCACCTCGGATAGGAAGCCGG<br>TGGCGGGAAGTTCGAAAATTGTTGA<br>GCTGAAATGTTTGCTCAAAAGCGCC | - |
| *ctl-2(ok1137)* | - | TACCCAGAAGCGTAATCCACAG<br>CATCTTGTCTGGCGAGAACTCG | - |
